# Vacuole phase-partitioning boosts mitochondria activity and cell lifespan through an inter-organelle lipid pipeline

**DOI:** 10.1101/2021.04.11.439383

**Authors:** AY Seo, F Sarkleti, I Budin, C Chang, C King, S-D Kohlwein, P Sengupta, J Lippincott-Schwartz

## Abstract

Functional linkage between mitochondria and lysosomes is crucial for survival under starvation and lifespan extension. Despite such co-dependency, the supportive pathways connecting mitochondria and lysosomes remain unclear. Here, we identify an inter-organelle lipid trafficking pathway linking yeast vacuole and mitochondria that results in increased mitochondria growth and respiratory activity under glucose starvation. The pathway depends on vacuolar phase-separated, lipid domains, which provide zones for: activation of the vacuolar proton pump; lipid droplet (LD) docking and internalization; and, lipid transfer from vacuole-to-ER-to-mitochondria. Partitioned vacuolar domains form through a specialized type of macro-autophagy, triggered only under acute glucose starvation, that delivers sterol-rich, endosomal-derived lipids to the vacuole. To balance this lipid influx, the vacuole reroutes lipids back to the ER to support both LD biogenesis and mitochondria growth and activity. Energy produced by enhanced mitochondrial activity then feeds back to support the inter-organelle lipid trafficking pathways to ensure survival under nutrient stress.

## INTRODUCTION

All cells experience periods of nutrient depletion and employ evolutionarily conserved pathways to effectively deal with this challenge. Two organelles, the lysosome and mitochondria, are particularly critical for cell survival under starvation. The lysosome breaks down cytoplasmic and membrane components acquired through bulk- and targeted-autophagy pathways, and the mitochondria metabolizes lipids for ATP production (Goldberg et al., 2009; Green et al., 2011; Seo et al., 2010; Yen and Klionsky, 2008; Zechner et al., 2012). In accomplishing these tasks, both organelles undergo changes: lysosomes remodel their surfaces to facilitate autophagic uptake and degradation, and mitochondria boost their membrane potential for effective oxidative phosphorylation. Deterioration in either set of responses compromises the cell’s survival under starvation and initiates age-related cellular defects.

Recent work in the budding yeast, *Saccharomyces cerevisiae*, has begun to shed light on how lysosomal and mitochondrial activities are coordinated to permit survival under starvation and lifespan extension. It has been shown, for example, that when starved cells sense an acute depletion of glucose they trigger uptake of lipid droplets (LDs) into the yeast’s lysosome-like vacuole in the process of micro-autophagy (Moeller and Thomson, 1979b; Tsuji et al., 2017; Seo et al., 2017; van Zutphen et al., 2014; Wang et al., 2014). Lipases within the vacuole interior break down the LDs into fatty acids, which, after being β-oxidized, are used by mitochondria as an alternative to glucose for ATP production (Weber et al., 2020). Disrupting various steps in this pathway significantly reduces yeast lifespan under starvation. Functional linkage between lysosomes and mitochondria has also been revealed in studies showing that increased vacuolar pH correlates with mitochondria dysfunction and cell lifespan reduction (Hughes and Gottschling, 2012) through cysteine toxicity (Hughes et al., 2020). Other studies have similarly shown lysosome function to be closely associated with mitochondrial decline in several age-related and metabolic disorders (Goodrum et al., 2019; Hughes et al., 2016).

To further clarify the vacuole’s role as key regulator of mitochondria function in starved cells, we focus here on the vacuole’s membrane properties. Prior work has shown that during nutrient stress, sterol-enriched microdomains similar to those found in giant unilamellar vesicles arise on the vacuole’s surface (Moeller et al., 1981; Moeller and Thomson, 1979a; Rayermann et al., 2017; Toulmay and Prinz, 2013; Tsuji et al., 2017), and that the integrity of these domains directly affects LD uptake into the vacuole (Seo et al., 2017; Wang et al., 2014). The membrane microdomains have been shown to arise from the demixing of a single liquid phase into two co-existing liquid phases, which can demix and remix. Domain formation is thought to be linked to availability of lipids and sterols that are delivered to the vacuole (Murley et al., 2015). Specific proteins localize within the different domains, helping to regulate or drive cooperative reactions by increasing the concentration of proteins and lipids while excluding other factors.

Despite these findings, it remains unclear where lipids/sterol forming the domains come from, whether the domains are necessary for the vacuole’s role in regulating mitochondria activity, and if so, what mitochondrial functions the domains support. Here, we addressed these questions in *Saccharomyces cerevisiae* by showing that phase separated microdomains arise by autophagic delivery of sterol-enriched endosomal membranes. Once formed, the domains permit more effective vacuolar acidification for substrate degradation (which is essential for avoiding mitochondrial dysfunction) and initiate a lipid pipeline for delivery of lipids into pathways that support mitochondria respiration/growth by supplying metabolic precursors and membrane components. Energy for driving this system depends on feedback from mitochondria, with impaired mitochondria leading to disappearance of the vacuole domains, blockage of the lipid pipeline, and early cell death. Mitochondria and vacuole health under nutrient stress are thus co-dependent on vacuole-centered, lipid trafficking pathways, with defects in these pathways likely underlying age-related mitochondria deterioration and vacuole dysfunction.

## RESULTS

### Mitochondrial growth and lifespan extension in starved cells depend on formation of vacuole sterol-enriched microdomains

Previous studies have shown that long-term survival under starvation in budding yeast occurs in response to acute reduction in glucose, as this triggers bulk autophagy and the targeted LD autophagy needed for cellular energy metabolism through mitochondria in the absence of glucose (Seo et al., 2017; Weber et al., 2020). Depleting glucose more slowly through gradual glucose restriction (i.e., gradual GR) only triggers bulk autophagy without targeted LD autophagy, hence cells do not survive long-term (Seo et al., 2017). Consistent with these findings, we found that mitochondria in cells experiencing 24 h of acute glucose restriction (i.e., acute GR) became extensively elongated and tubulated, occupying up to ~3 times more surface area compared to mitochondria in cells that were fed (Figure 1A). Cells experiencing the same period of gradual GR failed to show this change in mitochondrial morphology, with the mitochondria instead appearing small and fragmented (Figure 1A), consistent with their producing less energy (Scheckhuber et al., 2007). Examining cell lifespan under the different treatments revealed cells undergoing gradual GR could not survive long-term, whereas cells experiencing acute GR, with elongated/proliferated mitochondria, lived for weeks (Figure 1B).

**Figure 1.**
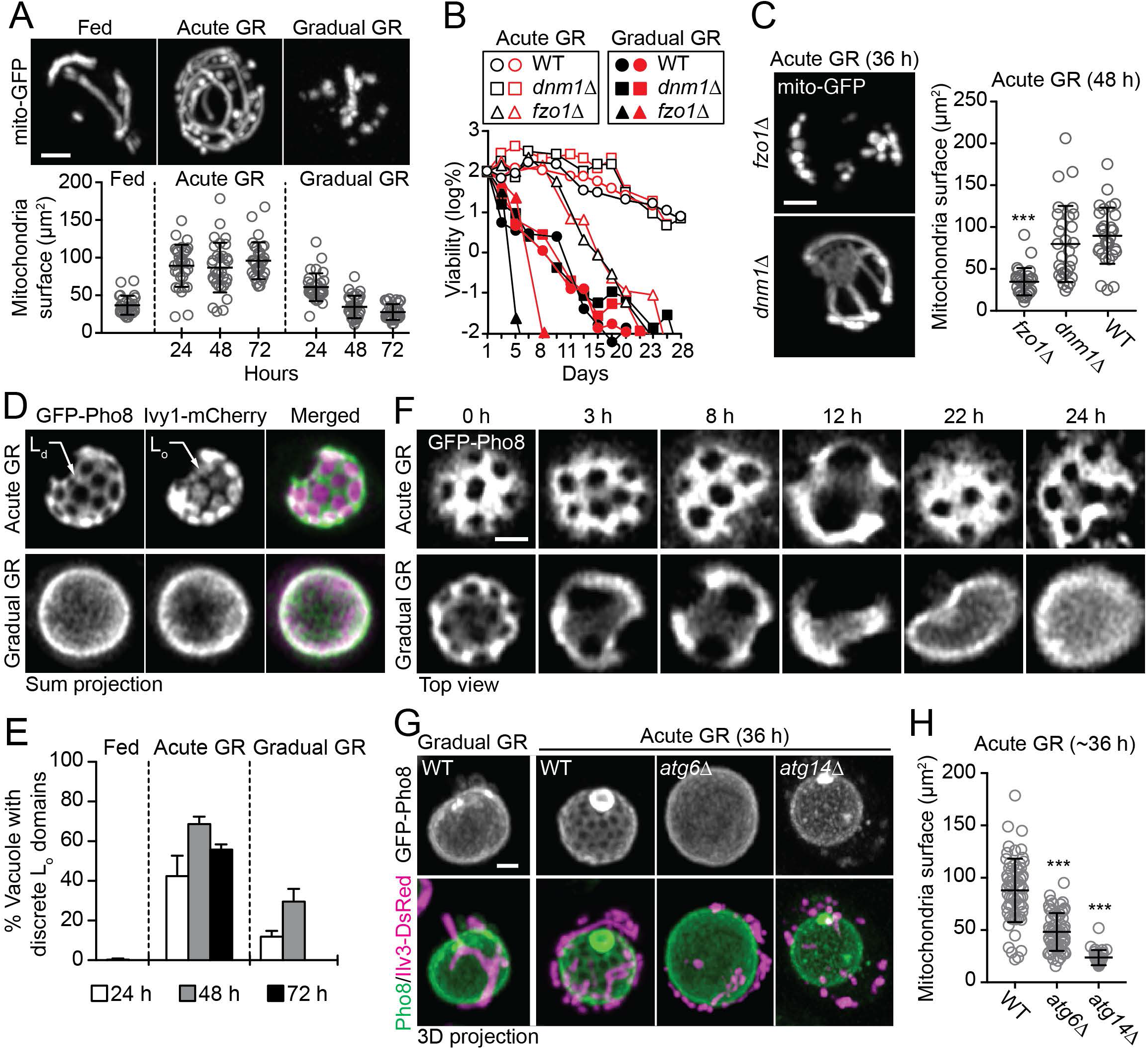
Mitochondrial growth and lifespan extension depend on vacuole microdomain formation. (A) Representative images at day 2 (*upper*) and quantification (*bottom*) of mitochondrial localized GFP (mito-GFP) from indicated conditions (n=~30 cells). (B) Cell survival data plotted as the log of a percentage viable cell number at day 1. Open and close symbols indicate independent experiments. (C) Representative images (*right*) and quantification (*left*) of mito-GFP structures in indicated deletion strains under acute GR (n=30 cells). (D) Representative images of cells expressing GFP-Pho8 and Ivy1-mCherry in indicated conditions. L_o_, liquid-ordered; L_d_, liquid-disordered. (F) Time-lapse analysis of cells expressing GFP-Pho8. (E) Quantification of vacuole L_o_ domain formation in cells in (D) (n=3-9; ≥100 cells/conditions). (G) Representative images of cells harboring GFP-Pho8 and Ilv3-DsRed. (H) Quantification of mito-GFP structures in indicated strains (n≥30 cells). Data are means ± SD. Size bars, 1μm. ANOVA with post-hoc Tukey HSD test: ***<0.001 vs WT.

To assess whether the survival advantage of cells experiencing acute GR was dependent on mitochondria membranes becoming highly fused and more abundant, we compared cells either lacking mitochondrial fusion activity (i.e., *fzo1Δ* cells)(Hermann et al., 1998) or lacking mitochondria fission activity (i.e., *dnm1Δ* cells)(Otsuga et al., 1998) under acute GR. The *fzo1Δ* cells could not tubulate and fuse their membranes under acute GR, so their mitochondria appeared small and fragmented (Figure 1C). This correlated with significantly reduced viability of the *fzo1Δ* cells relative to wild-type (WT) cells undergoing acute GR (Figure 1B). On the other hand, *dnm1Δ* cells lacking mitochondria fission activity, which have tubulated mitochondria, lived long-term during acute GR and exhibited highly tubulated/proliferated mitochondria, similar to that seen in control cells undergoing acute GR (Figure 1B, C). Together, the results indicated that long-term survival of cells under acute GR requires mitochondria to proliferate and become highly tubulated, which likely helps boost mitochondrial energy production.

We next examined whether any properties of the vacuole membrane could be important for enabling mitochondria to tubulate/proliferate under acute GR. Vacuole membranes in starved cells are known to form phase-separated liquid-ordered (Lo) microdomains enriched in sterol surrounded by a continuous liquid-disordered (Ld) phase that is less sterol enriched. Two probes shown to differentially distribute within these domains are Ivy1-mCherry, a vacuole protein known to reside in the sterol-rich, L_o_ membranes, and GFP-Pho8, a transmembrane vacuole protein that segregates into the sterol-poor, L_d_ membranes (Seo et al., 2017; Toulmay and Prinz, 2013; Wang et al., 2014). Co-labeling of cells with these two probes and starving either by acute or gradual GR revealed significant differences in the pattern of distribution of these markers (Figure 1D). Discrete vacuolar L_o_ microdomains containing Ivy1-mCherry were surrounded by a continuous L_d_ phase labeled by GFP-Pho8 in cells starved by acute GR. By contrast, in most cells starved by gradual GR, GFP-Pho8 and Ivy1-mCherry distributed homogenously across the vacuole surface (Figure 1D).

Quantification of the extent of L_o_ microdomain formation over different time periods of acute versus gradual GR revealed that while L_o_ microdomains formed at early times of gradual GR (24-48 h) in a small fraction of cells, they were no longer present in vacuoles at later time points (72 h) (Figure 1E). This contrasted with cells undergoing acute GR, in which the domains were maintained throughout 72 hours of starvation. No such discrete L_o_ microdomains were observed in vacuoles from well-fed cells (Figure 1E). Time-lapse imaging of a single vacuole expressing GFP-Pho8 over a 24 h period after starvation by either acute or gradual GR further revealed that the L_o_ microdomains excluding GFP-Pho8 were highly dynamic, undergoing merging, separation and disappearance (Figure 1F). Only in cells experiencing acute GR, however, did the dynamic L_o_ microdomains continue to appear over long periods (Figure 1F).

To explore whether vacuole membrane partitioning into L_o_ and L_d_ phases is required for mitochondrial elongation/proliferation in cells experiencing acute GR, we examined mitochondria morphology in cells in which vacuole phase partitioning was inhibited. Our prior work showed that vacuole partitioning into L_o_ and L_d_ phases in response to acute GR requires the autophagy proteins Atg6p or Atg14p (Seo et al., 2017; Wang et al., 2014). Examining cells lacking these proteins (i.e., *atg6Δ* or *atg14Δ* cells) under acute GR revealed that not only were there no distinct vacuole L_o_ and L_d_ phases (visualized using GFP-Pho8) but mitochondria (labeled with IIv3-mCherry) became highly fragmented and had reduced surface area (Figures 1G and 1H). Additionally, these cells did not survive long-term under acute GR (Seo et al., 2017). Therefore, preventing vacuole phase-partitioning in cells experiencing acute GR both inhibits mitochondria tubulation/proliferation and prevents long-term cell survival.

These results suggested that in cells undergoing acute GR, there are functional relationships involving phase-partitioned vacuole microdomains, mitochondria tubulation/proliferation, and cell survival. We aimed to clarify these relationships in the experiments described below; first, by examining how phase-partitioned vacuoles are generated under acute GR, and then by addressing what mitochondrial functions are served by these domains that can lead to extended cell lifespans.

### Ergosterol regulates the phase-partitioning characteristics of the vacuole membrane in cells undergoing acute GR

Phase-partitioned, L_o_ microdomains on the vacuole surface of starved cells are sterol-enriched (Toulmay and Prinz 2013), so it is widely believed these microdomains arise from increased sterol delivery to the vacuole (Tsuji et al., 2017). To directly test this idea, we controlled levels of ergosterol (the predominant sterol in yeast) in cells by replacing the promoter region of *ERG9* (encoding squalene synthase that provides the precursor for ergosterol) with the CYC1 promoter under the control of tetracycline-responsive tetO_2_ elements (Figures 2A, B). Upon doxycycline treatment, the *tetO_2_-ERG9* cells decreased ergosterol levels within 24 h of acute GR (Figure 2C). Examining the formation of L_o_ microdomains in these cells using the L_d_ marker, GFP-Pho8 revealed that discrete L_o_ microdomains did not form in doxycycline-treated cells undergoing acute GR; indeed, the percentage of cells with discrete L_o_ microdomains decreased in a doxycycline-dependent manner (Figure 2C-E). This suggested increased ergosterol is needed for L_o_ microdomain formation in the vacuole of cells starved acutely of glucose.

**Figure 2.**
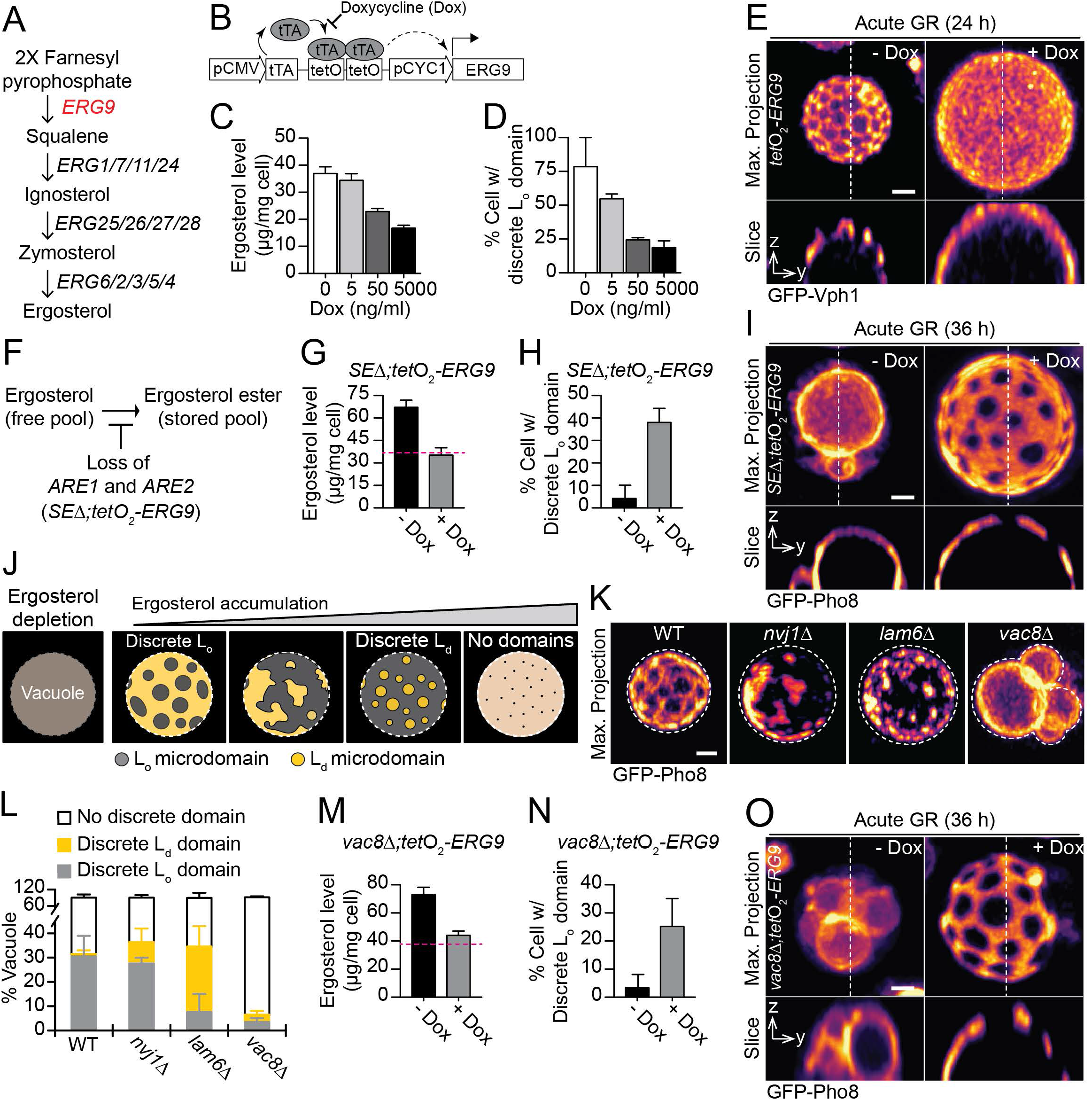
Ergosterol controls the phase-partitioning characteristics of vacuole membrane in cells undergoing acute GR. (A) Ergosterol synthesis pathway in *S. cerevisiae*. (B) Illustration of *ERG9* promotor region in the ergosterol repressible strain, *W303a;tetO_2_-ERG9*. (C) Quantification of ergosterol levels by Gas Chromatography-Mass Spectrometry (GC-MS) from *W303a;tetO_2_-ERG9* (n=3). (D) Quantification of *W303a;tetO_2_-ERG9* expressing GFP-Vph1 and displaying vacuole L_o_ domains (n=3; ~100 cells/condition). (E) Representative images of *W303a;tetO_2_-ERG9* cells expressing GFP-Vph1 under acute GR ± 5000ng/ml doxycycline (Dox). (F) Ergosterol esterification pathway in *S. cerevisiae*. (G) Quantification of ergosterol levels by GC-MS in the indicated strain ± 5000ng/ml Dox (n=4). (H) Quantification of the indicated strain that expresses GFP-Pho8 and display vacuole L_o_ domains during acute GR ± 5000ng/ml Dox (n=3; ~100 cells). (I) Representative images of cells in (H). (J) Illustration of vacuole phase-partitioning patterns depending on ergosterol levels. (K) GFP-Pho8 images from indicated deletion strains upon 24 h of glucose restriction. Dashed lines indicate vacuole boundaries. (L) Quantification of cells in (K) (n=3; ~80 cells). (M) Quantification of ergosterol levels by GC-MS in the indicated strain ± 5000ng/ml Dox (n=3). (N) Quantification of the indicated strain that expresses GFP-Pho8 and displays vacuole L_o_ domains during acute GR ± 5000ng/ml Dox (n=3; ~100 cells). (O) Representative images of cells in (N). Data are means ± SD. Size bars, 1μm. See also Movie S1.

To determine whether vacuole L_o_ domain development could also be inhibited by unusually high ergosterol levels, we increased those levels in cells undergoing acute GR by deleting the genes involved in ergosterol ester (SE) synthesis (i.e., *ARE1* and *ARE2*), which will block sterol ester formation and thereby increase free sterol levels (Zweytick et al., 2000). *ARE1* and *ARE2* deletions (i.e., *SEΔ*) were introduced into a *tetO_2_-ERG9* strain so we could additionally test the effect of lowering ergosterol levels by doxycycline treatment (Figure 2F). The *SEΔ;tetO_2_-ERG9* cells so generated displayed elevated ergosterol levels after 24 h of acute GR compared to ergosterol levels in similarly treated, control cells (Figure 2G, dashed line indicates ergosterol levels from WT cells in Figure 2C). Introduction of the L_d_ domain-preferring probe GFP-Pho8 revealed virtually no phase-partitioned vacuole L_o_ domains in these cells (Figure 2H), with GFP-Pho8 now homogenously distributed in the vacuole membrane (Figure 2I, -Dox). This suggested that vacuole ergosterol levels had risen so high in the *SEΔ;tetO_2_-ERG9* cells that discrete lipid phases could no longer be sustained. To confirm this, we lowered ergosterol levels in the cells by doxycycline addition and found that L_o_ domains reappeared (Figures 2G-I, +Dox).

The above results suggested that the phase-partitioning characteristics of the vacuole membrane in cells undergoing acute GR are linked to ergosterol levels-with low ergosterol levels producing a vacuole with no domains, medium ergosterol levels producing a vacuole with discrete L_o_ microdomains surrounded by a L_d_ phase, higher ergosterol levels producing a vacuole with L_d_ microdomains surrounded by a continuous L_o_ phase, and even higher ergosterol levels resulting in a vacuole with no microdomains (Figure 2J). As cells with vacuoles lacking discrete L_o_ microdomains do not survive longterm under acute GR (Seo et al., 2017), there must be exquisite regulation of the pathways for ergosterol delivery to and from the vacuole under acute GR. We examined these pathways as described in the sections below.

### Ergosterol pools necessary for producing L_o_ vacuolar domains come from a non-ER source

To investigate what membrane source supplies the needed ergosterol levels for producing discrete vacuolar L_o_ microdomains in vacuoles of cells starved by acute GR, we first focused on the possibility of ER. Extensive non-vesicular lipid trafficking occurs between the vacuole and ER. Both Lam6p and Nvj1p are ER proteins localized at ER-vacuole contact sites where they help regulate non-vesicular lipid trafficking between vacuole and ER through specific interactions with the vacuolar tethering protein Vac8p (Murley et al., 2015). We therefore deleted Lam6p or Nvj1p in GFP-Pho8-expressing cells to see if this prevented the vacuole from partitioning its membranes due to lack of sufficient ergosterol import by disruption of ER-to-vacuole lipid trafficking. It did not. We found not only that the vacuole did not lose its L_o_ phase in *nvj1Δ* and *lam6Δ* cells undergoing acute GR but that in some cells there appeared to be an overabundance of ergosterol in the vacuole because the L_d_-preferring protein GFP-Pho8 was now surrounded by a continuous L_o_ phase (Figure 2K, L). Vacuoles having discrete L_d_ microdomains surrounded by a continuous L_o_ phase were especially apparent in *lam6Δ* cells undergoing acute GR (Figure 2L and Movie S1). This suggested that inhibiting ER-vacuole lipid trafficking in cells undergoing acute GR actually leads to ergosterol accumulation on the vacuole surface rather than ergosterol depletion.

To further test this possibility, we examined the vacuole in cells lacking Vac8p, a lipid transfer protein that functions at all vacuole-ER contact sites. Again, we found that *vac8Δ* cells undergoing acute GR appeared to accumulate excess ergosterol in their vacuoles rather than experiencing ergosterol depletion. The excess ergosterol, in turn, led to the disappearance of L_o_ microdomains, with GFP-Pho8 becoming uniformly distributed across vacuole membranes (Figure 2K, L). Evidence that this was due to ergosterol overaccumulation in the vacuolar membrane (which, as shown above, can cause vacuole L_o_ domains to disappear, see Figure 2J), came from experiments in *vac8Δ;tetO_2_-ERG9* cells, which had *ERG9* under the control of the *tet*O_2_ promoter for lowering ergosterol levels through doxycycline treatment (Figure 2M; dashed line indicates ergosterol levels from WT cells in Figure 2C). Upon doxycycline-induced lowering of ergosterol levels in *vac8Δ;tetO_2_-ERG9* cells, L_o_ microdomains reappeared (Figure 2M-O). Together, these findings indicated that the ER does not supply ergosterol to the vacuole in cells undergoing acute GR.

### Autophagosomes deliver ergosterol to the vacuole during acute GR

Besides ER-vacuole contact sites, there are four other plausible routes that could be used for accumulating sterols and lipids in the vacuole to direct the formation of vacuolar L_o_ microdomains. These include 1) from LDs (Wang et al., 2014), 2) through intermembrane sterol transfer (Murley et al., 2015), 3) by vacuole sterol retention from multivesicular bodies (Tsuji et al., 2017), or 4) by autophagic delivery of sterol-rich membranes. To determine which of these routes was most responsible, we engineered GFP-Pho8-expressing cells with deletions in key proteins underlying each of these routes and then examined whether the vacuole formed L_o_ microdomains on its surface after 36 hours of acute GR.

In WT cells, as shown before, acute GR significantly increased the number of vacuoles containing L_o_ microdomains (~2.5x) compared to vacuoles undergoing gradual GR (Figure 3A). Cells lacking LD-sterol ester hydrolases to prevent ergosterol release from LDs (i.e., *tgl1Δ, yeh1Δ*, and *yeh2Δ* cells) or those deprived of ER-localized sterol transporters to block inter-membrane sterol transfer (i.e., *lam4Δ, lam5Δ*, and *lam6Δ* cells) showed no diminishment in capacity to form L_o_ microdomains under acute GR (Figure 3A). L_o_ microdomain formation under acute GR was only slightly diminished in cells lacking the sterol transporter proteins (i.e., Ncr1p or Npc2p) (Figure 3A). Based on these results, we concluded that ergosterol pools derived from routes 1-3 above are not major contributors to vacuole L_o_ microdomain formation under acute GR.

**Figure 3.**
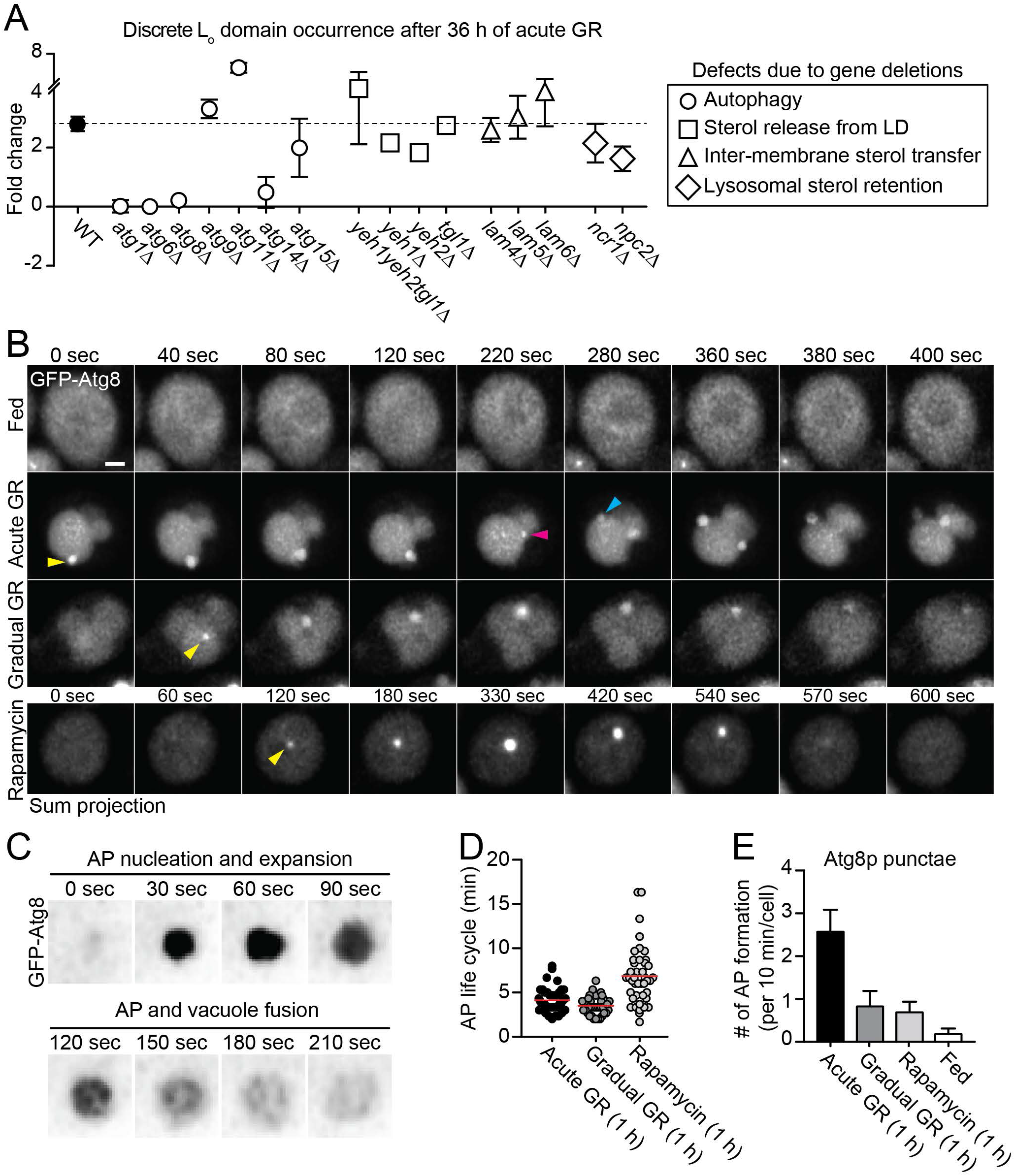
Acute GR enhances autophagosomal ergosterol delivery to the vacuole. (A) Quantification of relative increase of L_o_-containing vacuoles in indicated strains harboring GFP-Pho8 upon acute GR vs gradual GR (n=3; ~100 cells). (B) Time-lapse analysis of cells expressing GFP-Atg8 after 1h of acute or gradual GR, and 1h of 250nM rapamycin treatment. Colored arrowheads indicate distinct autophagosome (AP) formation events. (C) Time-lapse images of AP development in cells in (B). (D) Average timelines of AP formation in cells in (B) (n=~40; ~100 cells). (E) Quantification of AP numbers formed during 10 min in cells in (B) (n=~100 cells). Data are means ± SD. Size bars, 1μm. See also Figure S1 and Movie S2.

We therefore focused on whether autophagic membrane delivery is important for vacuole L_o_ microdomain formation. Supporting this possibility, deletion of genes encoding several autophagy components significantly reduced vacuole L_o_ microdomain development in cells starved acutely of glucose (Figure 3A). These included genes encoding proteins involved in pre-autophagosomal structure (PAS) formation (*ATG1* and *ATG8*) and those involved in autophagosome expansion (*ATG6* and *ATG14*), shown also in Figure 1G. Without these genes, cells did not form or sustain L_o_ microdomains on the vacuole surface in response to acute GR. Not all autophagy genes were equally important, however. Autophagy components that controlled membrane retrieval from either mitochondria or cytoplasmic vesicles (i.e., Atg9p) (Reggiori et al., 2005; Yamamoto et al., 2012) and the cytoplasm-to-vacuole (Cvt) targeting pathway (i.e., Atg11p) (Kim et al., 2001) were dispensable for L_o_ microdomain induction by acute GR (Figure 3A). Moreover, the major vacuolar lipase, Atg15p (Teter et al., 2001) was not required for vacuole L_o_ microdomain formation during early periods of acute GR. Together, these results indicated that the increased ergosterol levels in vacuolar membranes needed for forming L_o_ microdomains under acute GR is principally derived from autophagy, but only from particular autophagic processes.

### Acute GR boosts autophagic delivery to the vacuole

To better understand the role of autophagy in vacuole L_o_ microdomain development, we visualized autophagosome-vacuole fusion events in cells expressing GFP-Atg8p under four conditions: no GR, acute GR, gradual GR and rapamycin-treatment. Of these four regimens, only acute GR gave rise to long-lived L_o_ microdomains on the vacuole. Imaging of autophagosome-vacuole fusion events under these four conditions was performed after one hour of treatment, which was necessary to avoid buildup of GFP-Atg8 signal in the vacuole to levels that made imaging of fusion events too difficult. To trigger autophagy in this short time frame, cells were first grown in synthetic complete media with 2% glucose (i.e., Fed) to prevent autophagy induction. In order to induce autophagy, cells were then shifted to synthetic media without nitrogen sources and containing either 0.4% glucose (i.e., acute GR) or 2% glucose (i.e., gradual GR). Alternatively, they were treated with rapamycin.

Cells started nucleating GFP-Atg8p-positive autophagic puncta near the vacuole within 30 minutes of starvation by acute GR, gradual GR or rapamycin-treatment, but no autophagic puncta formed in well-fed cells (Figure 3B). Individual GFP-Atg8 puncta on the vacuole reached their maximal brightness within 90 sec and then disappeared over the next 2 min (Figure 3C), likely due to fusion with the vacuole membrane. This timecourse for docking and fusion of Atg8-labeled autophagosomes was similar whether autophagy was induced by acute GR or gradual GR and was slightly longer in cells treated with rapamycin (Figure 3D). Notably, significantly more autophagosome docking and fusion events occurred in cells undergoing acute GR (Figure 3B). Indeed, over a 10 min period, up to 3.7 times more GFP-Atg8p autophagosomes formed under acute GR compared to gradual GR, or rapamycin-treatment (Figure 3E; Movie S2). Furthermore, at any time-point, cells undergoing acute GR had a much larger number of autophagosomes docked on their surface (Figure S1A). Importantly, cells harboring both GFP-Atg8 and Atg6-yeRFP (which together serve as active PAS markers; Suzuki et al., 2001) showed increased Atg8-puncta that co-localized with Atg6p during 24 h of acute GR (Figure S1B).

These results indicated that acute GR significantly boosts autophagosome production and delivery to the vacuole relative to that from other starvation conditions or by rapamycin treatment. As long-term, vacuolar L_o_ microdomains only arise under acute GR, the data suggested that it is the heightened autophagic delivery under acute GR that provides the needed ergosterol for forming vacuolar L_o_ microdomains.

### Autophagic structures derived from endosomal membranes gives rise to the heightened autophagic activity under acute GR

We next investigated what membrane source gives rise to the autophagosomes undergoing increased rates of formation and fusion with the vacuole under acute GR. One potential source we examined was ER exit sites (ERESs), shown previously to be involved in autophagosome biogenesis (Graef et al., 2013). However, imaging cells after 2 h of acute GR versus gradual GR revealed no apparent difference in the spatial association between the ERES component, Sec16p-GFP and newly forming Atg8p puncta (Figure S2).

We then tested whether endosomes could give rise to the increased levels of autophagosomal structures seen forming at and fusing with the vacuole surface during 24 h of acute GR. Supporting this possibility, we found that most of the Atg8p-containing puncta appearing at the vacuole surface in response to acute GR were interacting with endocytic compartments labeled with the early/late endosome marker, Vps8-GFP (Figure 4A, C) or the recycling endosome/TGN marker, Sec7-GFP (Day et al., 2018) (Figure 4B, D). In addition, a GFP-FYVE construct, which labels PI3P-enriched membrane domains on endosomes (Burd and Emr, 1998), could be seen associating with the newly forming, expanding autophagosomes in time-lapse movies of cells after 1 h of acute GR (Figure 4E). Much less GFP-FYVE co-localized with newly emerging RFP-Atg8-labeled autophagosomes in rapamycin-treated cells or cells undergoing gradual GR (Figure 4F-H). Thus, the increased numbers of autophagosomes delivered to the vacuole under acute GR appeared to originate from a PI3P-enriched, endosomal source. As endosomes are ergosterol rich (Zinser et al., 1993), this led us to anticipate that endosome-derived autophagosomes that fuse with the vacuole under acute GR could be an important pathway for supplying membrane sterols to the vacuole membrane, enabling the formation of L_o_ microdomains.

**Figure 4.**
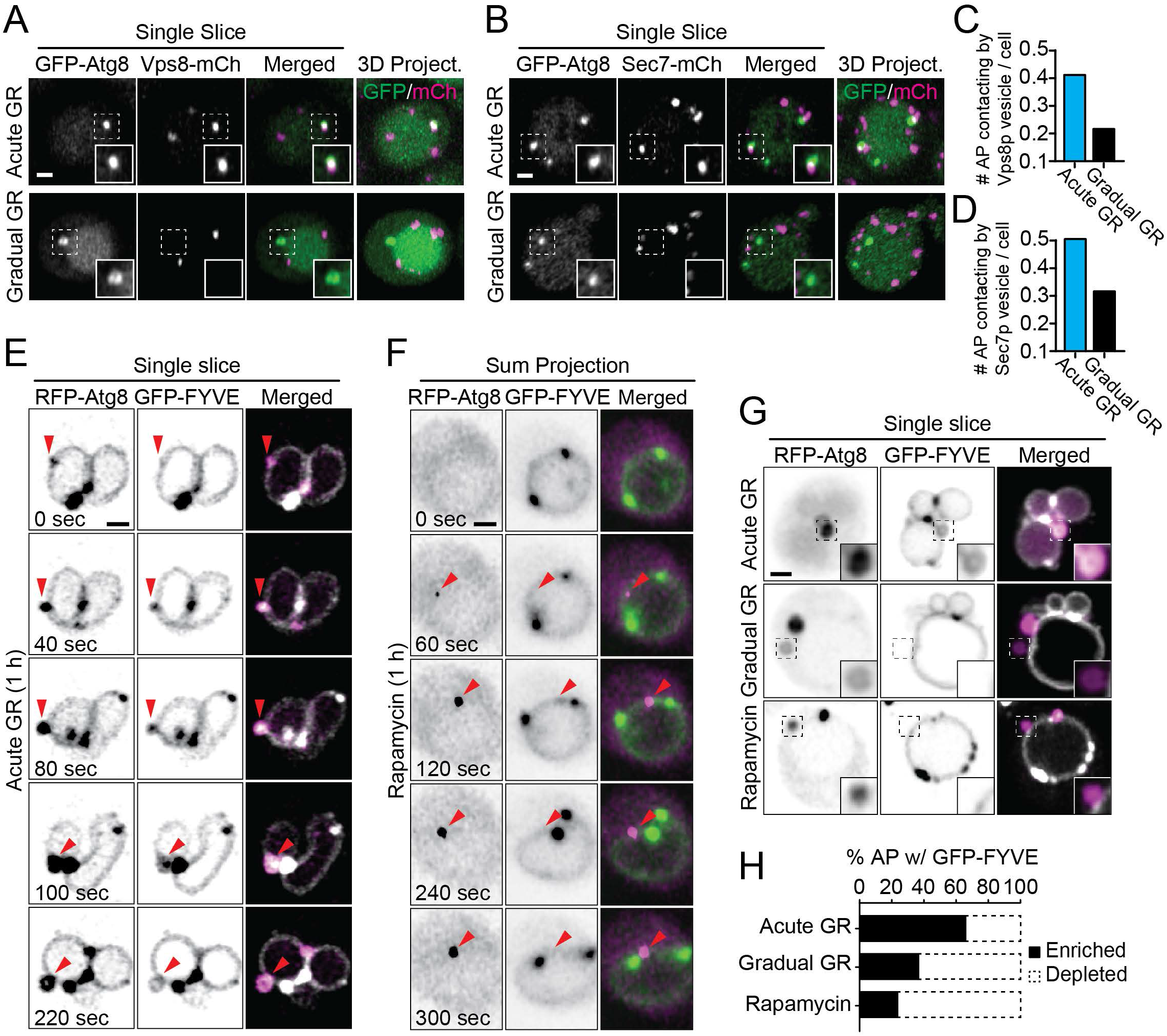
Enhanced autophagy during acute GR is supported by endosomal-derived autophagosomes. (A) Representative images of cells expressing GFP-Atg8 and Vps8-mCherry after 30 h of acute or gradual GR. (B) Representative images of cells expressing GFP-Atg8 and Sec7-mCherry after 30 h of acute or gradual GR. (C) Quantification of Atg8 puncta contacting with Vps8 vesicles in cells in (A) (n=~50). (D) Quantification of Atg8 puncta contacting with Sec7 vesicles in cells in (B) (n=~60). (E and F) Time-lapse images of cells expressing RFP-Atg8 and GFP-FYVE under indicated conditions. (G) Representative images of cells expressing RFP-Atg8 and GFP-FYVE after 3 h of acute GR, gradual GR, or 250nM rapamycin treatment. (H) Quantification of RFP-Atg8 puncta enriched by GFP-FYVE in cells in (G) (n=~50). Arrowheads indicate developing APs. Size bars, 1μm. See also Figure S2.

### Vacuolar L_o_ microdomains are internalized with docked LDs during acute GR

Prior work has shown that LDs dock exclusively at L_o_ microdomains on the vacuole surface under acute GR, with Atg14p helping to stabilize them (Seo et al., 2017; Wang et al., 2014) (Figure 5A and Movie S2). To further characterize the fate of LDs and vacuole membranes during this process, we performed time-lapse imaging of LDs bound to the vacuole in cells undergoing acute GR. Docked LDs on vacuole L_o_ microdomains were found to keep this surface association for several minutes before being internalized into the vacuole (Figure 5B). Further analysis of this process revealed that the LDs carried L_o_ membrane components, including sterol and proteins, when they internalized (Figure 5C). This was demonstrated in cells harboring both the L_o_ domain marker, Ivy1-mCherry, and the L_d_ domain marker, GFP-Pho8. In these cells, Ivy1-mCherry could be seen inside the vacuole as intact vesicles with no GFP-Pho8 signal (Figure 5C, See arrow heads; Figure 5D and Movie S3). A kymograph from a line drawn across the vacuole surface revealed that L_o_ domains exhibited localized motion over tens of seconds before suddenly disappearing from view (Figure 5E), presumably corresponding to their internalization into the vacuole interior.

**Figure 5.**
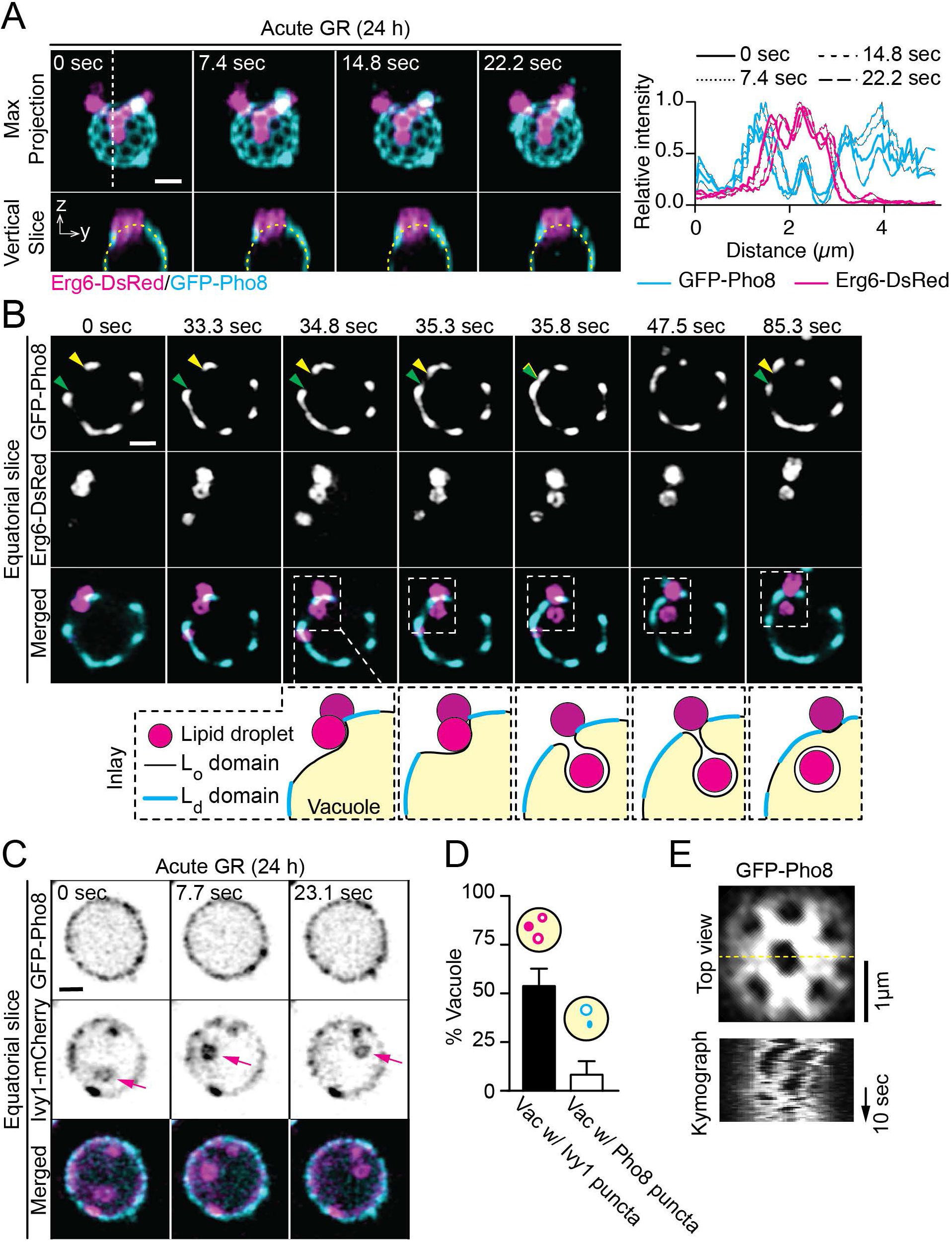
Vacuolar L_o_ microdomains are selectively internalized with LDs. (A and B) Representative time-lapse images of cells expressing Erg6-DsRed and GFP-Pho8 (*right*). Fluorescence intensity plots along the yellow dashed lines in the bottom y-z slices (*left*). (B) Time-lapse images of cells expressing GFP-Atg8 and Erg6-DsRed after 36 h of acute GR with scaled illustrations of the region indicated in the top row. Arrow heads indicate L_o_/L_d_ membrane boundaries. (C) Time-lapse images of cells expressing GFP-Pho8 and Ivy1-mCherry. Arrows indicate internalized Ivy1-decorated vesicles in the vacuole. (D) Quantification of cells containing vacuolar Ivy1 or Pho8 puncta in (C) (n=3; ~70 cells). (E) Kymograph analysis along the yellow dashed line in the top panel from cells expressing GFP-Pho8 after 24 h of acute GR. Size bars, 1μm. See also Movie S3.

Together, these results indicated that in cells undergoing acute GR, vacuole L_o_ microdomains are continuously and selectively internalized into the vacuole. As sterols in L_o_ domains are also internalized during this process, there must be continued input of new sterols to the vacuole to prevent it from losing all its L_o_ domains. A likely possibility was that such sterol input derived from the endosome-associated autophagy described above, which was triggered by acute GR.

### Vacuole phase partitioning leads to increased vacuole acidity

LDs internalized into the vacuole only release their stored lipids when pH-sensitive vacuole lipases are active to digest the LDs. We tested whether the vacuole L_o_ domains play a role in facilitating this turnover by examining the vacuole distribution and activity of Vph1p, the Vo component of the vacuole proton pump (i.e., V-ATPase), which acidifies the vacuole by delivering protons into it (Manolson et al., 1992). Using Vac8-GFP to label L_o_ domains (Peng et al., 2006; Subramanian et al., 2006), we found that in response to acute GR, Vph1p partitioned to the L_d_ phase surrounding L_o_ domains on the vacuole surface (Figure 6A), as previously reported (Toulmay and Prinz, 2013; Tsuji et al., 2017). Interestingly, using cells in which Vph1p was endogenously tagged with RFP, we observed significantly more Vph1p being recruited to and/or stabilized at the vacuole surface (labeled with Vac8-GFP) under acute GR (Figures 6B and 6C), suggesting possible greater proton pump activity under these conditions.

**Figure 6.**
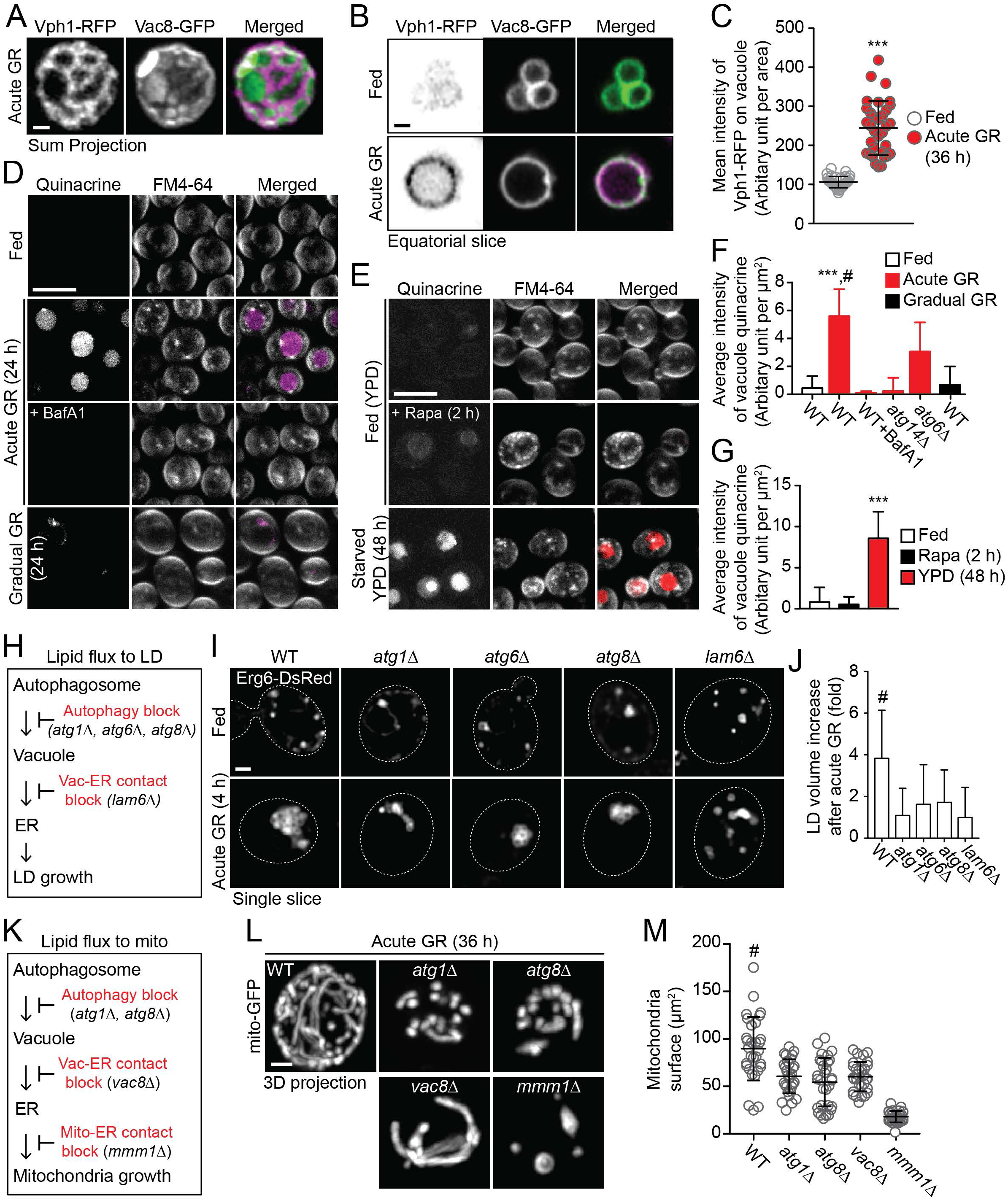
Vacuole phase partitioning leads to vacuole acidification, helping LD and mitochondrial growths during acute GR. (A and B) Representative vacuole images of cells in which Vph1p was endogenously tagged with RFP and that harboring Vac8-GFP. (C) Quantification of Vph1-RFP recruitment on the vacuole surface in cells in (B) (n≥30). (D and E) Confocal images of cells pulsed by quinacrine and FM4-64 from indicated conditions ± 5μM Bafilomycin A1 (BafA1), or 250nM rapamycin (Rapa). (F and G) Quantification of vacuolar quinacrine signals from cells in (D, E, and S3A; n=~100). (H) Proposed inter-organellar lipid pipeline to lipid droplet (LD). (I) Representative images of LD (labeled by Erg6-DsRed) in indicated strains under untreated (Fed) or 4 h of acute GR. Dashed lines indicate cell boundaries. (J) Quantification of relative LD volume change in cells from (B) upon 4 h of acute GR compared to the LD volume in the untreated (Fed) cells (n=44-74). (K) Proposed inter-organellar lipid pipeline to mitochondria (mito). (L) Representative images of indicated strains expressing mito-GFP. (M) Quantification of mito-GFP structures in cells in (L) (n=30). Size bars, 10μm [D and E] and 1μm [A, B, I and L]. ANOVA with post-hoc Tukey HSD test: ***<0.001 vs Fed, #<0.001 vs deletion strains. See also Figure S3.

To test this, we measured vacuole pH levels using the acid sensitive dye quinacrine. This lipophilic cationic compound accumulates in the vacuole when its lumen is acidic (Preston et al., 1989). Quinacrine in the vacuole was much greater in cells undergoing acute GR than in cells that were unable to phase-partition their vacuoles (i.e., cells undergoing gradual GR) or in well-fed cells (Figure 6D, F). Bafilomycin A1, a vacuolar-type H^+^-ATPase inhibitor completely prevented vacuolar quinacrine accumulation in cells undergoing acute GR (Figure 6D, F), confirming that the dye accumulation reflected V-ATPase activity in these cells.

We further examined the acidity of vacuoles in cells incubated in YPD medium. Here, rapamycin treatment, which does not give rise to significant L_o_ microdomain formation (Seo et al., 2017; Toulmay and Prinz, 2013), showed no vacuole quinacrine accumulation, similar to untreated cells (Figure 6E, G). However, cells incubated in YPD medium that were starved for 48 h, accumulated large amounts of quinacrine in their vacuole. Notably, these cells had partitioned vacuoles with L_o_ microdomains (Figure S3B). This suggested that vacuoles with L_o_ microdomains may have acidic lumens. Supporting this idea, *atg14Δ* or *atg6Δ* cells under acute GR, in which no vacuole L_o_ microdomains are formed, were found to have significantly diminished quinacrine signals, indicating their lumens were not acidic (Figure 6F and S3A).

Together, these results indicated that partitioning of V-ATPase components to the vacuole L_d_ phase under acute GR promotes their assembly into an active proton pump, leading to increased acidification of the vacuole. The boosted vacuole acidity would then facilitate recycling and digestion of substrates (e.g., LDs) internalized into the vacuole during acute GR.

### Vacuole lipids are routed to ER for LD biogenesis under acute GR

To survive long-term under acute GR, cells must continually generate new LDs that can be routed to the vacuole for digestion into metabolites useable for mitochondrial activity. LDs originate from the ER, so lipids delivered into LDs are either newly synthesized in the ER or trafficked to the ER (Hariri et al., 2018). As lipid biosynthesis is inhibited under starvation, lipid trafficking pathways must perform this role. We reasoned, therefore, that continued input of lipids into the vacuole through autophagy during starvation could provide a source of lipids for trafficking back to ER (via vacuole-ER contact sites) for LD biogenesis and mitochondrial function.

To test this idea, we examined how blocking either autophagy or vacuole-ER lipid trafficking affects LD formation in cells undergoing acute GR (Figure 6H). Using Erg6-DsRed to label LDs, we quantified the number and volume of LDs in cells lacking the autophagy proteins, Atg1p, Atg6p or Atg8p, or lacking the vacuole-ER sterol transfer protein, Lam6p. Removal of any of these proteins was found to significantly reduce the amount of LDs being formed during 4 h of acute GR, with little or no impact on LDs in well fed cells (Figures 6I and 6J). These results thus supported the idea that vacuole lipids are routed back to ER for LD biogenesis in cells undergoing acute GR.

### Acute GR triggers a lipid pipeline from vacuole to mitochondria

Given our data indicating that lipids return to the ER from the vacuole in cells undergoing acute GR, we next asked whether some of these lipids are transferred to mitochondria (via ER-mitochondria contact sites) for expansion of its membranes, enabling greater energy production by mitochondria. To test this possibility, we examined the effect on the mitochondria of blocking different steps in the proposed lipid flux pathway, including from autophagosome to vacuole, from vacuole to ER, and from ER to mitochondria, using yeast strains with deletions in each transfer step (Figure 6K). Cells lacking *ATG1* or *ATG8* were used to block autophagosome biogenesis, while *vac8Δ* and *mmm1Δ* cells were respectively employed to block vacuole-to-ER transport (Pan et al., 2000) and ER-to-mitochondria lipid transfer (Kornmann et al., 2009). By analyzing mitochondrial structures in cells also harboring mito-GFP, we found that blocking any of these steps significantly perturbed mitochondria growth under acute GR as measured by their surface area (Figure 6L, M). This suggested the mitochondrial growth defect was, in fact, due to the inability of these cells to transfer lipids into the mitochondria.

Thus, together with our other findings, these results supported our proposal that under acute GR, a lipid pipeline initiated by autophagic lipid delivery to the vacuole triggers the formation of phase-partitioned vacuole domains that then enable trafficking of lipids back to the ER for LD formation and into the mitochondria to support mitochondrial growth.

### Mitochondrial ATP provides the energy for the lipid pipeline

The question still remained, however, what energy source could drive the various steps required by the lipid pipeline. During starvation, glycolysis is unavailable, leaving only the mitochondria’s production of ATP as an energy source. We therefore examined whether the bioenergetic state of mitochondria could underlie the functioning of the phase-separated, vacuolar microdomains. It is known that inhibiting mitochondrial fusion in *fzo1Δ* cells leads to mitochondrial energy loss, while inhibiting mitochondria fragmentation in *dmn1Δ* cells helps enhance mitochondria energy metabolism (Scheckhuber et al., 2007). We examined vacuoles using GFP-Pho8 in *fzo1Δ* and *dmn1Δ* cells undergoing acute GR and found that L_o_ microdomains were absent in *fzo1Δ* cells but present in *dmn1Δ* cells (Figure 7A). This suggested that mitochondrial energy production is needed for generating the phase-partitioned vacuole L_o_ domains.

**Figure 7.**
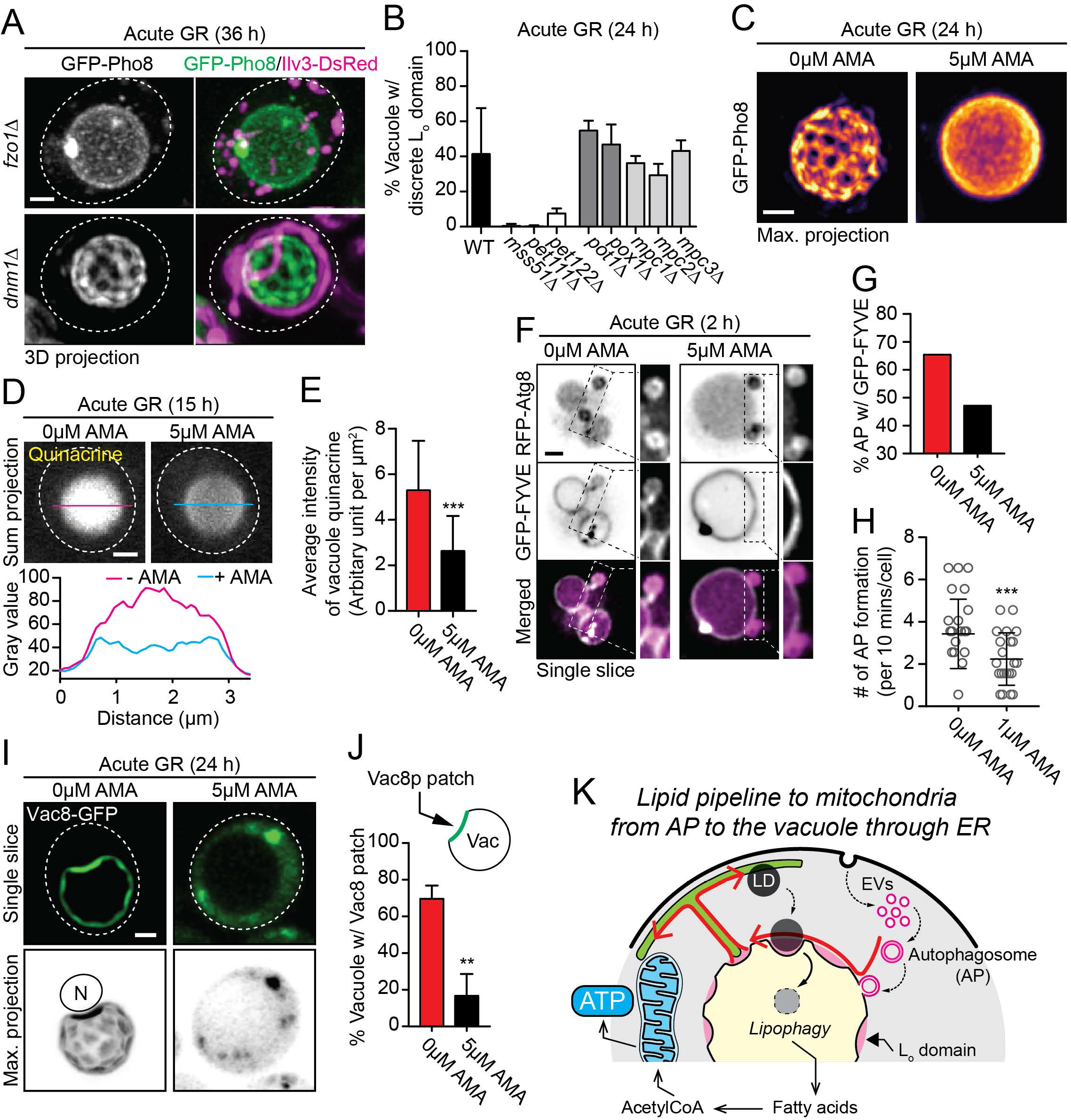
Mitochondria provide the energy for operating the inter-organellar lipid pipeline during acute GR. (A) Representative images of indicated deletion strains expressing GFP-Pho8 and Ilv3-DsRed. (B) Quantification of vacuole L_o_ domain formation in indicated deletion stains harboring GFP-Pho8 (n=3; ~100 cells). (C) Representative images of cells expressing GFP-Pho8 under acute GR ± 5μM Antimycin A (AMA). (D) Representative images of cells pulsed by quinacrine under acute GR ± 5μM AMA (*top*). Fluorescence intensity plots along the magenta and cyan lines (*bottom*). (E) Quantification of vacuolar quinacrine signals from cells in (D) (n=~140). (F) Representative images of cells expressing RFP-Atg8 and GFP-FYVE under acute GR ± 5 μM AMA. (G) Quantification of RFP-Atg8 puncta enriched by GFP-FYVE in cells in (F) (n=~120). (H) Quantification of autophagosome (AP) formation in cells harboring GFP-Atg8 under acute GR ± 1 μM AMA (n=~20 cells). (I) Representative images of cells expressing Vac8-GFP under acute GR ± 5μM AMA. N, nucleus. (J) Quantification of Vac8-GFP recruitment to the vacuole (vac) surface in cells in (I) (n=3; ~100 cells). (K) Illustration of proposed intracellular lipid pipeline (red arrows) that is operated upon acute GR and necessary for mitochondrial energy metabolism. EVs, endosomal vesicles. Size bars, 1μm. Dashed lines indicate cell boundaries. t-test: **<0.01, ***<0.001 vs 0μM AMA.

To test this idea further, we examined whether impairing other mitochondrial energy-related steps caused vacuole L_o_ microdomains (visualized using GFP-Pho8) to disappear in cells undergoing acute GR. Notably, perturbation of mitochondrial electron transfer chain (ETC) complexes in *mss51Δ, pet111Δ*, and *pet122Δ* cells (Decoster et al., 1990; Kloeckener-Gruissem et al., 1988; Poutre and Fox, 1987) led to the complete loss of vacuolar L_o_ microdomains upon 24 h of acute GR (Figure 7B). However, similarly treated cells without genes for peroxisomal β-oxidation (i.e., *POT1* and *POX1*) (Dmochowska et al., 1990; Mathieu et al., 1997) or mitochondrial pyruvate import (i.e., *MPC1, MPC2*, and *MPC3*) (Bricker et al., 2012) showed WT-like L_o_ microdomains in the vacuoles (Figure 7B). When we used antimycin A (AMA) as an alternative to gene deletions for inhibiting mitochondrial ETC activity and energy production, vacuolar L_o_ microdomains in cells undergoing acute GR also disappeared (Figure 7C). These results suggested that energy production through the mitochondrial ETC is essential for vacuole L_o_ microdomain formation.

Given the finding that vacuole L_o_ microdomain formation was associated with increased vacuole acidity, we examined whether treating starved cells with AMA would block that activity. Indeed, we found that AMA treatment significantly reduced the extent that quinacrine was taken up by the vacuole during acute GR (Figure 7D, E). This indicated that vacuoles could not sustain their low pH under acute GR without mitochondrial energy production.

We next investigated what pathways associated with vacuole L_o_ microdomain formation/maintenance were dependent on mitochondrial energy production. First, we looked at macroautophagy-mediated delivery of the endosomal-derived sterol-enriched membranes to the vacuole. Previous work has shown AMA treatment reduces total autophagosome production (Adachi et al., 2017; Graef and Nunnari, 2011). Examining cells co-expressing RFP-Atg8 and GFP-FYVE, we found that AMA treatment preferentially reduced the production of autophagosomes labeled by GFP-FYVE in acutely glucose restricted cells (Figure 7F-H). These results suggested that mitochondrial-derived ATP production is necessary for formation and/or delivery of endosome-derived autophagosomes (enriched in PI3P binding sites) to the vacuole.

Another vacuole-related pathway we considered for evaluating the role of mitochondria energy flux was vacuole-to-ER lipid trafficking. To test this possibility, we examined AMA’s impact on the distribution of the vacuole-ER contact site protein, Vac8p. In untreated cells undergoing acute GR, Vac8p was localized on the vacuole surface (Figure 7I), with enrichment in L_o_ microdomains and particularly at vacuole-nuclear envelope contact sites (Subramanian et al., 2006). Upon ATP-reduction with AMA-treatment, Vac8p dissociated from vacuolar membranes and redistributed into the cytoplasm, where it was either diffusely distributed or localized on large, punctate structures (Figures 7I and 7J). Thus, when mitochondrial ATP production is inhibited by AMA treatment in cells starved acutely of glucose, vacuole-to-ER contact sites disassemble and vacuole to ER lipid trafficking is potently disrupted.

Together, these results suggested that mitochondrial energy production feeds back to organize inter-organellar lipid trafficking pathways under nutrient stress, by facilitating and sustaining vacuole partitioning into L_o_ and L_d_ phases through induction of specialized autophagy of endosomal membranes. In turn, the functions of these phase partitioned domains, which include activation of the V-ATPase, stimulation of lipophagy and reorganization of Vac8p at vacuole-ER contact sites, are also dependent upon mitochondria energy production.

## DISCUSSION

The work here delineates a mechanism linking vacuole and mitochondrial functions under nutrient starvation centering on a lipid pipeline involving autophagosomes, vacuole, ER, LDs and mitochondria (Figure 7K). This inter-organelle system allows mitochondria to grow and be active for energy metabolism when glucose is scarce. As discussed below, the characteristics and regulators of the lipid pipeline have important implications for understanding and manipulating the effects of nutrient stress in health, disease, and aging.

We found that the primary integrator of the lipid pipeline was the vacuole’s phase partitioned membrane domains. Sterol-enriched L_o_ microdomains surrounded by a continuous L_d_ phase, described previously in several studies (Moeller et al., 1981; Toulmay and Prinz, 2013; Tsuji et al., 2017), were uniquely formed and sustained in response to acute glucose depletion. This vacuole phase-partitioning pattern only arose and was maintained when ergosterol generated through autophagy reached specific levels within the vacuole; too much or too little ergosterol and there was no L_o_ microdomain formation. The phase-partitioned domains performed three crucial functions necessary for mitochondria energy adaptation and long-term cell survival under starvation: 1) lipophagy, 2) vacuole pH reduction and 3) vacuole-ER contact formation. Preventing the vacuole from phase partitioning blocked all three functions, resulting in mitochondrial decline and early cell death.

The first of these functions, lipophagy, has been described before and involves vacuolar L_o_ microdomains providing sites for autophagic uptake of LDs into the lipase-rich vacuole interior (Moeller and Thomson, 1979b; Seo et al., 2017; Tsuji et al., 2017; Wang et al., 2014). This allows for breakdown of LD-stored triacylglycerols into free fatty acids, which get further processed into simpler substrates in peroxisomes for mitochondrial use in ATP production. We visualized LD uptake into the vacuole in live cells and found that L_o_ microdomains were internalized with LDs. This suggested that in order for L_o_ microdomains to be sustained long-term on the vacuole surface, new L_o_-domain-forming lipids must be delivered to the vacuole for replacing those internalized with LDs. Searching for the replacement pathway, we found a candidate in the form of a specialized type of macro-autophagy that delivered endosomal-related membranes to the vacuole only under acute GR. Using Vps8, Sec7 or FYVE probes to mark endosomes, we showed this endosomal-related autophagic pathway supplied the vacuole with new membranes at rates over 3 times that of other forms of macro-autophagy that continue to occur under these conditions. Endosomal membranes have high sterol levels, so we reasoned that the endosome-related autophagic pathway induced by acute GR supplied the needed sterols both for forming phase-partitioned L_o_ domains and replacing those internalized into the vacuole by lipophagy. Consistent with this interpretation, blocking upstream autophagic components in cells undergoing acute GR led to vacuole ergosterol reduction and disappearance of L_o_ microdomains.

Our evidence revealed a second function served by vacuole phase partitioning, to lower the vacuole’s pH. This would allow the vacuole to be more active for substrate degradation. We found the V-ATPase component Vph1p under acute GR was localized to L_d_ domains surrounding L_o_ microdomains, as also reported by others (Toulmay and Prinz, 2013; Tsuji et al., 2017), so we reasoned this localization might help concentrate and/or stabilize Vph1p with other V-ATPase components. This, in turn, might lead to the assembly and activation of the proton pump. Supporting this possibility, we found more Vph1p was vacuole-associated when the vacuole had partitioned its membranes into L_o_ and L_d_ domains, and the vacuole was more acidic. Importantly, blocking vacuole phase partitioning in cells undergoing acute GR prevented the increased vacuole acidification.

The third function served by vacuole phase partitioning, ER contact site formation, was shown by imaging the major vacuole-ER contact site protein Vac8p in cells undergoing acute GR. In these cells, Vac8p was enriched in vacuole L_o_ domains, particularly those in close contact with the nuclear envelope, where significant vacuole-ER lipid exchange occurs (Hariri et al., 2018; Murley et al., 2015). Deleting Vac8p in cells undergoing acute GR led to massive ergosterol accumulation (and loss of vacuole domain formation), suggesting the function of Vac8p activity at vacuole-ER contact sites is to alleviate excess vacuole lipids by transferring them into the ER. Delivery of the vacuole lipids into the ER, in turn, could facilitate cell survival under acute GR by providing lipids for new LD biogenesis and for routing lipids to mitochondria for membrane proliferation (Stewart and Yaffe, 1991). Supporting this latter idea, we found that either deleting Vac8p to disrupt vacuole-to-ER lipid trafficking or removing ER-mitochondria contact proteins to disrupt ER-to-mitochondria lipid trafficking led to the creation of small, fragmented mitochondria and/or reduced LD biogenesis, with cells unable to survive long-term under acute GR.

These results pointed to vacuole phase partitioning as a critical component of an inter-organelle lipid pipeline that serves to boost mitochondrial growth and activity in starved cells (Figure 7K). The pipeline takes lipids from phase-partitioned vacuoles and directs them into the ER, from which the lipids move into either LDs or mitochondria. As a consequence, LDs are continually formed, and mitochondria continuously supplied with lipids for expansion of their membranes. But what membrane pathway would provide the lipids for running this pipeline? Our findings indicate it is the endosome-related autophagic pathway induced under acute GR. Without induction of this autophagic pathway, not only were phase-partitioned vacuole domains not maintained and lipophagy inhibited, but vacuole acidification was reduced, LD formation diminished, and mitochondrial growth impeded.

To understand what energy source could drive the various steps required by the lipid pipeline described above, we turned to mitochondria ATP production, as glycolysis is unavailable under acute GR. Inhibiting mitochondrial oxidative phosphorylation using AMA blocked the entire lipid pipeline, resulting in no vacuole phase-partitioning, no vacuole acidification, reduced LD degradation, and mitochondria becoming fragmented and small. Two steps in the lipid pipeline were specifically inhibited by blocking mitochondrial ATP production. One was the formation of endosome-related autophagosomes; the other, vacuole-ER lipid trafficking. The latter resulted from Vac8p becoming dissociated from vacuole membranes by AMA treatment. Because of Vac8p’s central role in vacuole-ER lipid transport, this caused the shut-down of lipid trafficking into the ER from the vacuole. With no influx of lipids, the ER in AMA-treated cells could no longer supply mitochondria with new membranes to proliferate, resulting in mitochondria fragmentation and size diminishment.

Our findings that mitochondria energy production is necessary for the different steps of the lipid pipeline induced under acute GR is relevant for understanding how various diseases involving defective mitochondria and vacuoles could imperil cells under nutrient stress. Cells continuously experience intervals of nutrient depletion as they age and develop. Our data suggests that if mitochondria become less capable of oxidative phosphorylation, the resulting shutdown of compensating pathways to rebuild active mitochondria will make cells less likely to survive nutrient stress. Our findings also suggest that one way to revitalize mitochondria in cells where mitochondrial energy production has become unreliable is to starve the cells acutely of glucose. Based on our findings, this would help rejuvenate the mitochondria by initiating the lipid pipeline for delivering vacuole-derived lipids obtained from autophagy to mitochondria via ER. Induction of the lipid pipeline by glucose starvation would also lead to increased acidification of the vacuole, making it more active for uptake of amino acids. This might help alleviate conditions of amino acid/cysteine toxicity arising from failed uptake of these substrates into the vacuole, recently shown to be associated with mitochondrial dysfunction and lifespan reduction in yeast (Hughes and Gottschling, 2012; Hughes et al., 2020). Given that lipid trafficking pathways, autophagy and lifespan extension by glucose starvation are conserved among many organisms, our findings in yeast could be applicable to other eukaryotes.

## STAR METHODS

Detailed methods, supplemental information with Key Resources Table are provided in the online version of this paper.

## ACKNOWLEDGMENTS

We thank Drs. Andreas Mayer, Christian Ungermann, Joel M Goodman, Tomohiro Yorimitsu, and William A Prinz for the kind gift of plasmids, and all members of the Lippincott-Schwartz laboratory for discussion. F.S. was supported by the DK Molecular Enzymology W901, funded by the Austrian Science Fund (FWF).

## AUTHOR CONTRIBUTION

A.Y.S. and J.L. designed experiments. A.Y.S. and F.S. performed all experiments. I.B. designed strains and performed ergosterol analysis. C.C., C.K., S. D. K., and P.S. provided materials. A.Y.S. analyzed data and prepared figures. A.Y.S. and J.L. wrote the manuscript. All authors provided critical review of the manuscript and approved the final version of the manuscript.

## SUPPLEMENTAL INFORMATION

### METHODS

#### Yeast Strains, Media, and Plasmids

Genetically manipulated *S. cerevisiae* strains and plasmids used in this study are listed in Tables S1 and S2, respectively. All chemicals used in this study were purchased from Sigma-Aldrich unless otherwise noted. Yeast media used in this study were prepared according to Seo et al. (Seo et al., 2017) with slight modification. Briefly, synthetic dextrose media were prepared with 0.17% (wt/vol) Difco yeast nitrogen base (without amino acids and ammonium sulfate) and 10 mM K2HPO4 with 0.5% (wt/vol) ammonium sulfate (SD) or without ammonium sulfate (SD-N) at 80% of the final volume. To prepare synthetic complete (SC) medium, 0.067% (wt/vol) Complete Supplement Mixture minus leucine and uracil (MP Biomedicals, Santa Ana, CA) was supplied to the above SD medium. Rich medium (YPD) was prepared with 1% (wt/vol) Bacto yeast extract, 2% (wt/vol) Bacto peptone, and 2% (wt/vol) dextrose with or without 1.6% (wt/vol) Bacto agar. All media were autoclaved at 121°C for 15 min before the supplement of appropriate auxotrophic nutrients. Supplemental auxotrophic nutrient and 20% (wt/vol) dextrose stocks were prepared according to Sherman et al. (Sherman, 2002) and were sterilized by using a 0.02 μm syringe filter. A final concentration of 250 μg/mL geneticin (G418) or 100 μg/mL nourseothricin (NAT) was provided when cells harbor kanMX or natMX cassettes, respectively. For synthetic media without nitrogen source, SD-N media containing 2% dextrose or 0.4% dextrose were prepared at pH 6.5. With sterile water, the final volume of each medium was adjusted to 100% before use.

To generate *pBS-ATG6-yeRFP-HIS3* or *pRS416-prCUP1-yeRFP-ATG8*, yeast-enhanced mRFP (yemRFP) was amplified from pRS415*-yemRFP* by PCR using primers: 5’ - atcgggatcctatggtttcaaaaggt--gaagaa & 3’- atcgtgtacatttatataattcatccataccacca, or 5’ - atcgactagtggatcctatggtttcaaaaggtgaagaa & 3’- atcgcccgggggtttatataattcatccataccacc, respectively. The resulting PCR products were used to replace GFP sequences in *pBS-ATG6-3×GFP-His3* using BamHI and BsrGI restriction sites, or in *pCuGFP-AUT7(416)* using SpeI and SmaI restriction sites. To generate *pRS415-prCPY-GFP-PHO8*, DNA fragment containing *prCPY-GFP-PHO8* sequence was amplified by primers: 5’- catctagacttctgcacaagaagccatattgacagagcag & 3’- gaggatccgagttagataggatcagttggtcaactcatgg. Using XbaI and BamHI restriction sites, the resulting product was subcloned into pRS415 vector. To construct *pRS413-prGPD-ILV3-DsRed*, ORF of *ILV3* was amplified from yeast genomic DNA (extracted from BY4742 strain by using Yeast DNA Extraction Kit (Thermo Fisher Scientific Inc., Waltham, MA)) with the following primers: 5’- tggaatccggatgggcttgttaacgaaagttgc & 3’- tctcgagtgagcagcatctaaaacacaaccg. Using Gateway system, the amplified *ILV3*-ORF without stop codon was inserted into a Gateway ‘entry’ vector using HpaII/MspI and XhoI restriction sites, and was further subcoloned to the ‘destination’ vectors as followed by the manufacture’s instruction (Thermo Fisher Scientific Inc., Waltham, MA).

#### Yeast Growth, Glucose Restriction, and Genetic Manipulations

All experiments were performed with biologically independent, colony-isolated strains. Unless otherwise noted, all strains were grown at 30°C. Glucose restriction (GR) experiments were performed according to Seo et al. (Seo et al., 2017). Briefly, yeast cells from frozen stocks were patched onto YPD agar plates. After 2 days of recovery, yeast patches were inoculated into 5 mL of YPD in a 14 mL polypropylene tube and were grown in a shaking incubator with 250 rpm. After ~18 h, the overnight (O/N) culture was diluted 1:100 in 5 mL of SC containing 2% dextrose plus leucine and uracil. After O/N, 0.05 ~ 0.1 OD unit (OD_600nm_ × mL) of yeast cells was transferred to 5 mL of SD media containing either 0.4% dextrose (acute GR) or 2% dextrose (gradual GR) plus appropriate auxotrophic nutrients and antibiotics. For long-term GR experiments, yeast cells remained in the aforementioned SD media without any nutrient replenishment. For shorter-term GR experiments (<4 h), SD-N media with only 0.4% dextrose (acute GR) or 2% dextrose (gradual GR) were used, unless otherwise noted. In order to avoid any starvation stress, yeast cells for all fed conditions in this study were grown under 20 mL of SC (or YPD) media with frequent nutrient replenishment by supplying fresh media, while maintaining OD_600nm_ around 0.5.

Genomic integrations for promotor replacement, knockout, and fluorescent protein tagging were performed through homologous recombination. To generate AYS1837, tagRFP-natMX cassette with homologous sequence at 5’ and 3’ ends was amplified from pIM700 with primers: 5’ -ataaagacatggaagtc gctgttgctagtgcaagctcttccgcttcaagcggagctggtgctggtatg & 3’ - tatttaatgaagtacttaaatgtttcgctttttttaaaagtcctcaaaatccggtagaggtgtggtca. Ergosterol titratable strain, tetO_2_-ERG9 were generated by PCR products from the template pCM224, as previously described (Degreif et al., 2019). AYS1824 was generated by two PCR products: one from the template pRS415 using primers: 5’ - gtggttgttcagcacggcttgcagcaagagcgccaaaacagattgcaagagaccgcagttaactgtggg & 3’ - ttgtatatctatcaagggcttgcgagggacacacgtggtatggtggcagtgagaatctttttaagc aaggattttc, and the other from the template pRS414 using primers: 5’ - aagtaaacagacacattacgttagcaaaagcaacaataacaaacacaaccgattgtactgagagtgcacc & 3’ - aaaatttactataaagatttaatagctccacagaacagttgcaggatgcccctatttcttagcatttttgacg. AYS1845 was generated by PCR products from the template pRS415 using the following primers: 5’ - atcctacgaagacaggaactgagcaaactataagggtgttctttcttctgtactatatatacatttgcaactatggaccgcagttaactgtggg & 3’ - ttctgagaagaaaattttgataaaaattataatgcctagtcccgcttttgaagaaaatcagagaatctttttaagcaaggattttc. AYS1837 was generated by using a linearized pBS-*ATG6-yeRFP-HIS3* (digested by KpnI). AYS1901 and AYS1903 were generated, as previously described (Day et al., 2018). Unless otherwise indicated, transformation of yeast cells was performed by using EZ-Yeast Transformation Kit (MP Biomedicals, Santa Ana, CA) with slight modification. Genomic integrations were routinely verified by colony PCR.

#### Cell Survival Measurements

Survivability of yeast cells under GR was routinely monitored by using a vital dye (i.e., erythrocin B) and was evaluated by measuring chronological life spans according to Alvers et al. (Alvers et al., 2009).

#### Drug Treatment

Stock solutions (1 mM) of rapamycin and bafilomycin A1 were prepared in dimethyl sulfoxide (DMSO) and were used with indicated final concentrations. Doxycycline stock (4 mg/mL) was prepared in water, filter-sterilized, and added to SD or SC media with indicated final concentrations. Antimycin A was dissolved in ethanol to make 1 mM stock solution and added to SD media with indicated final concentrations. Stock solutions for quinacrine hydrochloride (HCL) (40 mM) and FM4-64 (1 mM) were prepared in sterile water and DMSO, respectively. All stock solutions were stored at −20°C.

#### Microscopy

All imaging experiments were performed at 30°C in SD or SC media unless otherwise noted. Epifluorescence images of live cells were acquired at room temperature using EVOS FL Cell Imaging System (Thermo Fisher Scientific Inc., Waltham, MA) equipped with DAPI (357/44 Ex; 447/60 Em), GFP (470/22 Ex; 510/42 Em), and RFP (531/40 Ex; 593/40 Em) light cubes. Confocal fluorescence images of live cells were acquired using Zeiss LSM 880 microscopy with a 63×Plan-Apochromat 1.4 NA oil objective (Carl Zeiss Inc., Thornwood, NY) for quinacrine HCL (458 Ex; 508-643 Em) and for FM4-64 (514 Ex; 642-735 Em). Airyscan images of live or fixed cells expressing proteins tagged by GFP and/or RFP were acquired using Zeiss LSM 880 Airyscan microscopy with a 63×Plan-Apochromat 1.4 NA oil objective (Carl Zeiss Inc., Thornwood, NY). GFP and RPF fusion proteins were exited with 488 and 561 nm lines, respectively. For live cell imaging, 100 to 200 μL of cell cultures was directly placed to Nunc™Lab-Tek™ II Chambered Coverglass (Thermo Fisher Scientific Inc., Waltham, MA). In case of overcrowding live cell culture (>2.0 OD_600nm_), ~0.05 OD unit of cells from each culture condition was prepared with the corresponding cell-free supernatants and was placed to the above imaging chamber. For fixed cells, 20 μL of fixed samples was placed on agarose pads under a 22×22 mm coverglass and imaged at room temperature. Airyscan raw data was first processed via the commercial Zen software package (Carl Zeiss Inc., Thornwood, NY) and was further analyzed by ImageJ (National institutes of Health, Bethesda, MD) with indicated plugins.

#### Yeast Fixation for Imaging

Cells were fixed using cold 4% paraformaldehyde with 3.4% sucrose at room temperature for 15 min. Then, cells were washed twice with KPO_4_/Sorbitol (0.1 M potassium phosphate with 1.2 M sorbitol at pH 7.5). Afterwards, cell pellets were resuspended in 50 μL KPO_4_/Sorbitol with DAPI at 1 μg/mL concentration and were incubated at room temperature for 10 min. After washing twice with KPO_4_/Sorbitol, fixed cells were resuspended in the same buffer and stored at 4°C.

#### Ergosterol Manipulation and Analysis

Ergosterol titratable strains (tetO_2_-ERG9) grown under appropriate SD or SC media for 24 h in the presence of varying amounts of doxycycline were collected and washed once with sterile water by centrifugation. The collected cell pellets were stored at −80°C prior to ergosterol analysis. To extract and analyze ergosterol, 2 OD unit of cells alongside an internal cholesterol standard was used as previously described (Rodriguez et al., 2014). Analysis was performed by gas-chromatography mass spectrometry (GC-MS) on an Agilent 7890 GC and 5975 MS using a 30m DB-5 ms column in selected ion monitoring mode. The temperature of the GC oven was held at 80°C for 1 min, followed by a ramp of 20°C·min^-1^ to 280°C with a 20 min hold time, followed by a ramp of 20°C·min^-1^ to 300°C with a 2 min hold time. Quantification of ergosterol was performed using peak areas of the *m/z* 386 ion, which were normalized to extraction efficiency using the internal standard (cholesterol, *m/z* 396 ion), and external ergosterol standards run in parallel. Values were normalized to cell dry mass using a standard curve for the tested strain. Of note is that without doxycycline treatment, both tetO_2_-ERG9 cells and its parental strain (i.e., W303a) exhibited similar ergosterol levels.

#### Quinacrine Uptake Analysis

To evaluate vacuolar pH in stationary phase cells under SD or YPD media containing 2% dextrose or 0.4% dextrose, we assessed vacuolar quinacrine accumulation as previously described (Flannery et al., 2004) with slight modification. Briefly, after growing for 15 to 24 h in SD media or 48 h in YPD media, 2 OD unit of cells was transferred to 1 mL of Buffer A (YPD containing 0.4% dextrose, 100 mM HEPES (pH 7.5), and 200 μM quinacrine HCL) in a 14 mL polypropylene tube. After 25 min in a shaking incubator with 250 rpm, one microliter of FM4-64 stock solution (1.6 mM) was added to the above 14 mL tube. Followed by 5 min incubation in the same shaking incubator, cells were chilled on ice for 5 min and were quickly washed three times with Buffer B (0.2% dextrose plus 100 mM HEPES (pH 7.5) in sterile water) at room temperature. After 30 min from the beginning of the first washing step using Buffer B, cells were mounted on Nunc™Lab-Tek™ II Chambered Coverglass and were immediately imaged using the microscope method described above. All image data from different strains and growth conditions were collected within 30 min at room temperature with identical imaging acquisition settings. Mean intensity of quinacrine signals from the center of vacuole area (*r* = 0.6 μm) was measured with background signal subtraction.

#### Organelle Analysis

For analysis of mitochondrial surface area, Z-stack images of GFP-labeled mitochondria from different strains with varying growth conditions were collected using Zeiss LSM 880 Airyscan microscopy in a SR mode as described previously. Total mitochondrial surface area in a single yeast cell was estimated by the sum of individual mitochondrion’s surface area determined by the software plugin, Yeast_MitoMap (Vowinckel et al. 20015) in ImageJ. To determine vacuole localization of Vph1p, AYS1718 strains grown under fed or 36 h of acute GR conditions were imaged using Zeiss LSM 880 Airyscan microscopy in a SR mode as described previously. From a single Z slice image showing tagRFP-label vacuole(s) with maximum diameter, Vph1p-tagRFP intensity on the vacuole surface (V_SUR_INTENSITY_) was estimated: V_SUR_INTENSITY_ = (V_DENSITY_ - L_DENSITY_)/(V_AREA_ - L_AREA_), where V_DENSITY_ and L_DENSITY_ mean raw integrated density values from the areas of vacuole (V_AREA_) and vacuole lumen (L_AREA_), respectively. For LD volume analysis, cells harboring *pRS315-ERG6-DsRed* under fed or 4 h of acute GR conditions were chemically fixed and imaged using Zeiss LSM 880 Airyscan microscopy in a SR mode as described previously. Z-stack images of DsRed-labeled LDs from a single yeast cell were analyzed by the software plugin, 3D Objects Counter (Bolte and Cordelieres) in ImageJ to determine total LD volume in the cell (LD_Total_VOL_). In each deletion strain, average of LD_Total_VOL_ from the fed condition (i.e., LD_Aver_VOL_FED_) LD_Total_VOL_ after acute GR (i.e., LD_Total_VOL_AGR_) were calculated, and the fold increase of LD volume after acute GR was estimated by the ratio of LD_Total_VOL_AGR_ minus LD_Aver_VOL_FED_ to LD_Aver_VOL_FED_.

#### Statistical Analysis

Statistical analysis was performed using GraphPad Prism 7 software for ANOVA with post-hoc Tukey HSD test or Student’s t-test (GraphPad Software, San Diego, CA).

**Table S1.**
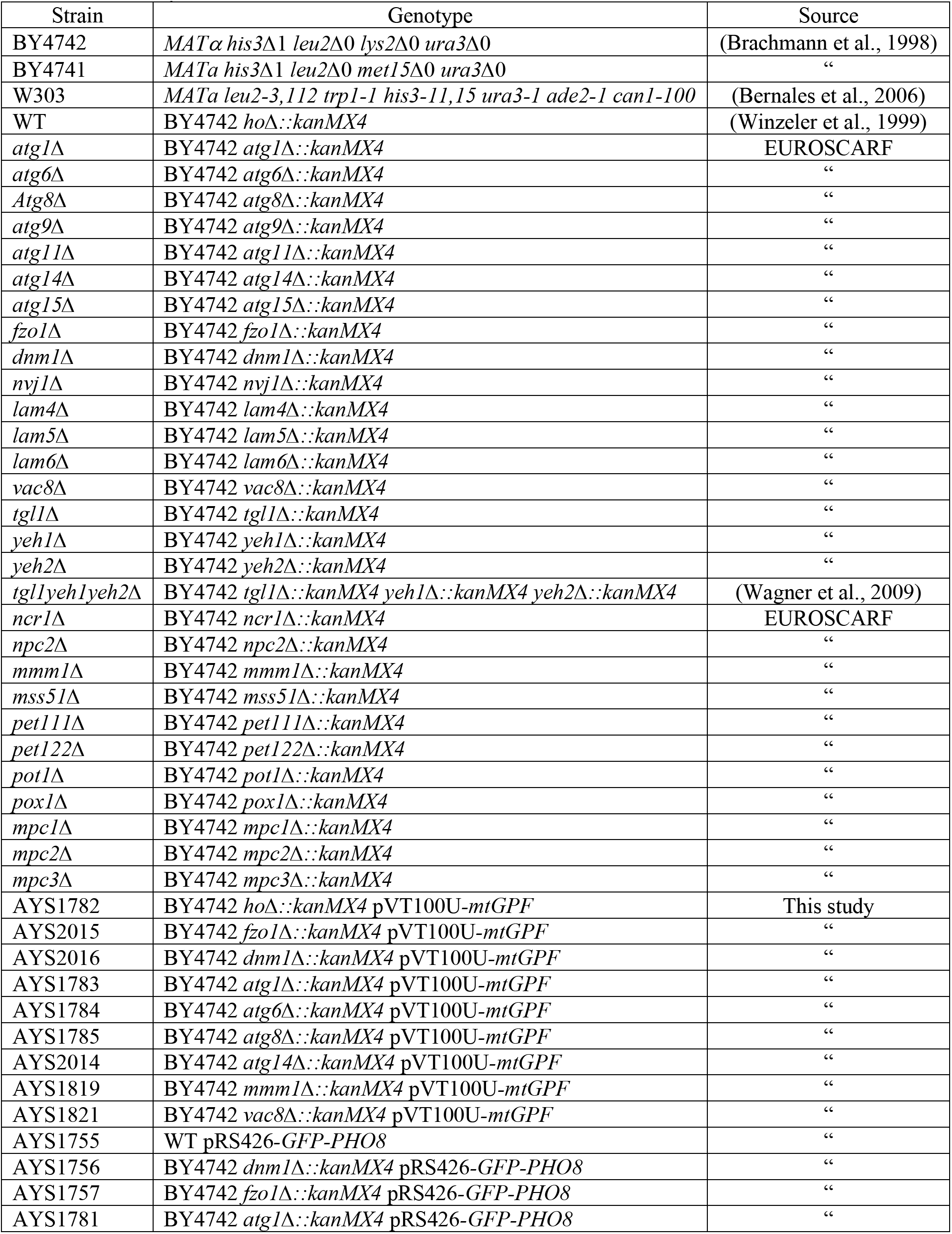
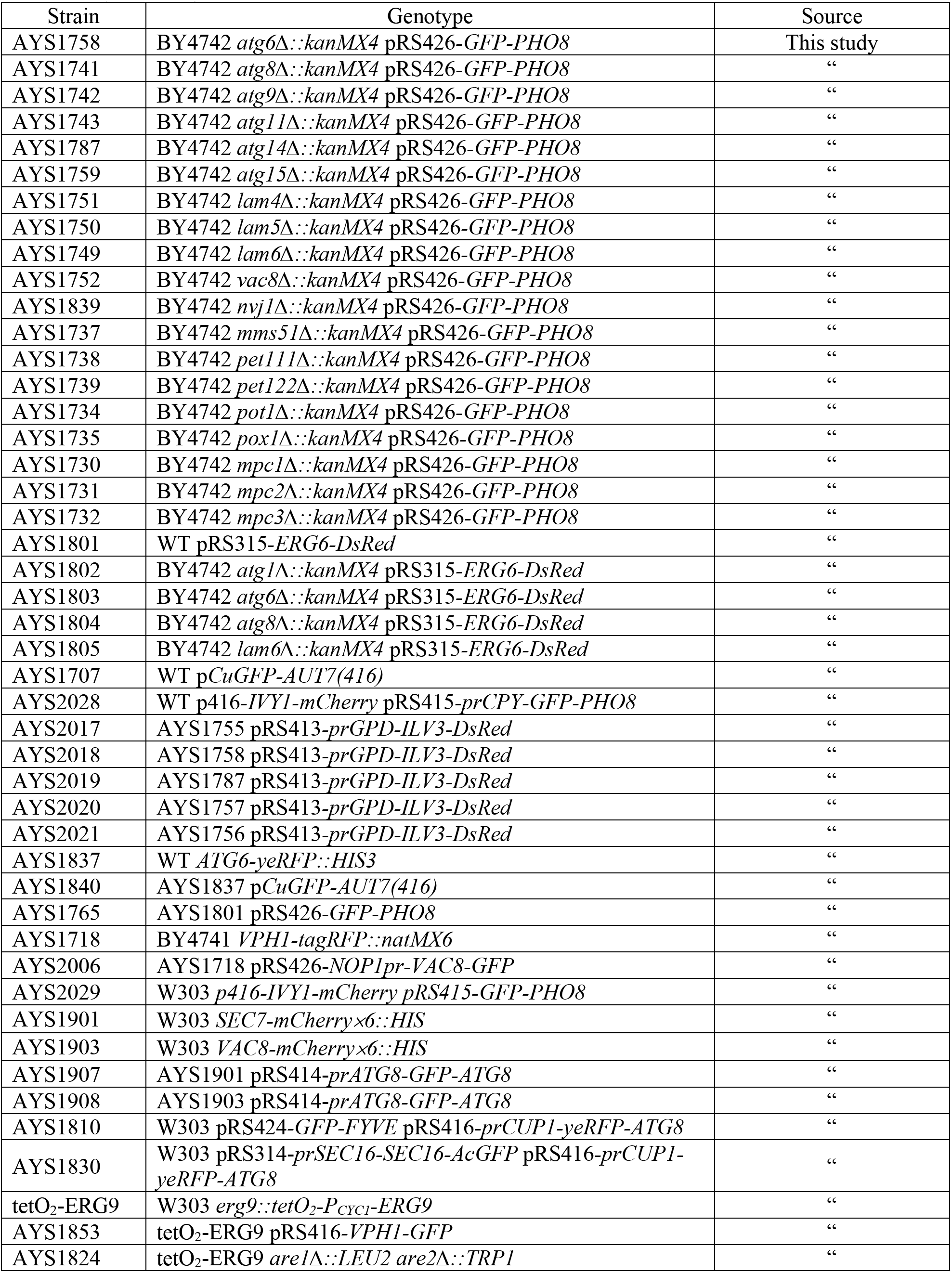
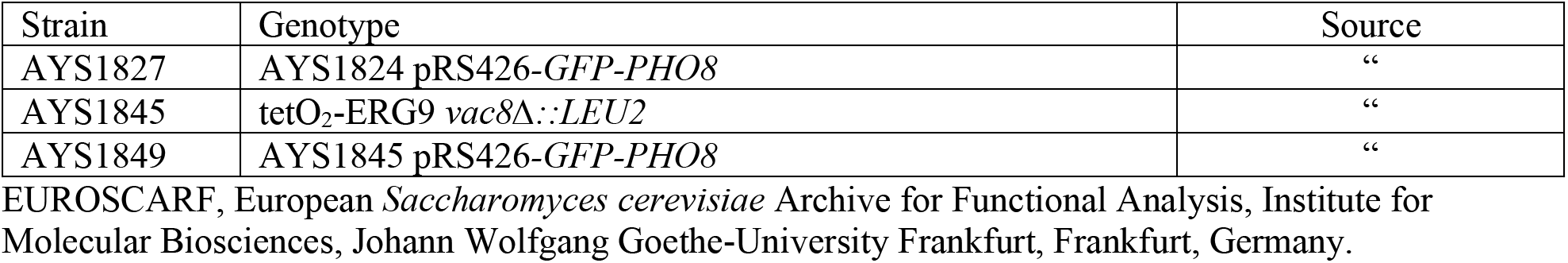
List of yeast strains, Related to STAR Methods

**Table S2.** List of plasmids, Related to STAR Methods

| Plasmid | Organelle monitored | Reference |
| --- | --- | --- |
| pCuGFP-AUT7(416) | Autophagosome | (Kim et al., 2001) |
| pRS414-*prATG8-GFP-ATG8* | “ | (Abeliovich et al., 2003) |
| pRS416-*prCUP1-yeRFP-ATG8* | “ | This study |
| pBS-*ATG6-yeRFP-HIS3* | “ | This study |
| pRS426-*NOP1pr-VAC8-GFP* | Vacuole | (Subramanian et al., 2006) |
| p416-*IVY1-mCherry* | “ | (Toulmay and Prinz, 2013) |
| pRS416-*VPH1-GFP* | “ | (Dawaliby and Mayer, 2010) |
| pRS426-*GFP-PHO8* | “ | “ |
| pRS415-*prCPY-GFP-PHO8* | “ | This study |
| YIplac211-*VPS8-mCherry2B×6* | Endosome | (Day et al., 2018) |
| YIplac211-*SEC7-mCherry2B×6* | “ | “ |
| pRS424-*GFP-FYVE* | “ | (Burd and Emr, 1998) |
| pRS314-*prSEC16-SEC16-AcGFP* | ER | (Yorimitsu and Sato, 2012) |
| pVT100U-*mtGFP* | Mitochondria | (Westermann and Neupert, 2000) |
| pRS413-*prGPD-ILV3-DsRed* | “ | This study |
| pRS315-*ERG6-DsRed* | LD | (Binns et al., 2006) |
| pBS-*ATG6-3×GFP-His3* | - | (Wang et al., 2014) |
| pIM700 | - | (Malcova et al., 2016) |

## SUPPLEMENTAL FIGURE LEGENDS

**Figure S1.**
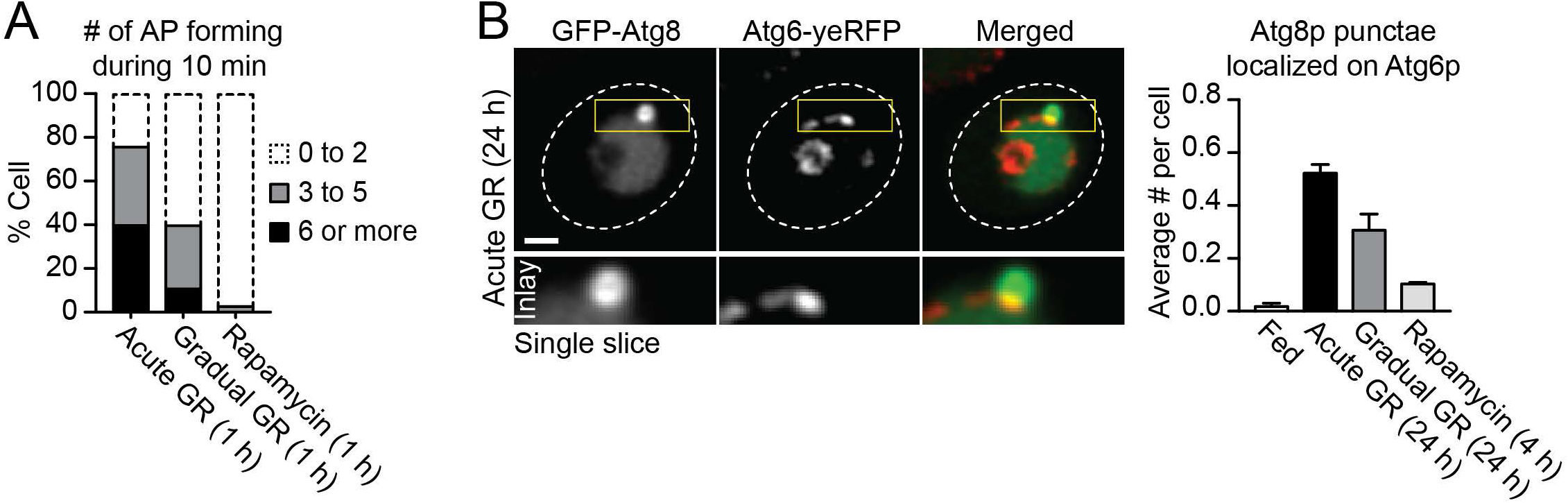
Acute GR enhances autophagosome biogenesis, related to Figure 3. (A) Quantification of cells expressing GFP-Atg8 with indicated numbers of Atg8 puncta formation over the course of 10 min upon ~3 h of acute or gradual GR, and 3 h of 250 nM rapamycin treatment (n=~70 cells/conditions). (B) Representative images of cells expressing GFP-Atg8 and Atg6-yeRFP under 24 h of acute GR (*right*). Quantification of GFP-Atg8 puncta localized to Atg6p-yeRFP structures in untreated (Fed), 24 h of acute or gradual GR, and 4 h of 250 nM rapamycin treatment (n=2; ~100 cells/conditions). Data are means ± SD. Size bars, 1μm.

**Figure S2.**
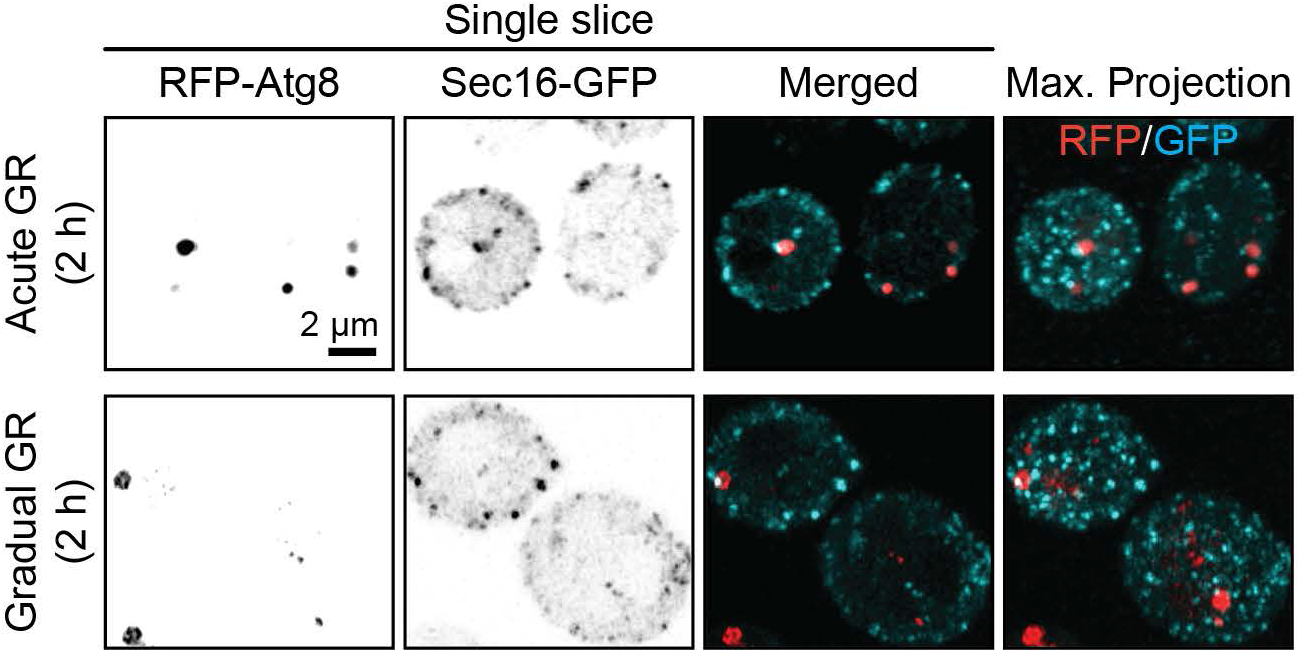
ER-exit site serves as membrane source for autophagosome biogenesis in both cells under acute GR and gradual GR, related to Figure 4. Representative images of cells expressing RFP-Atg8 and Sec16-GFP under 2 h of acute GR or gradual GR. Size bars, 1μm.

**Figure S3.**
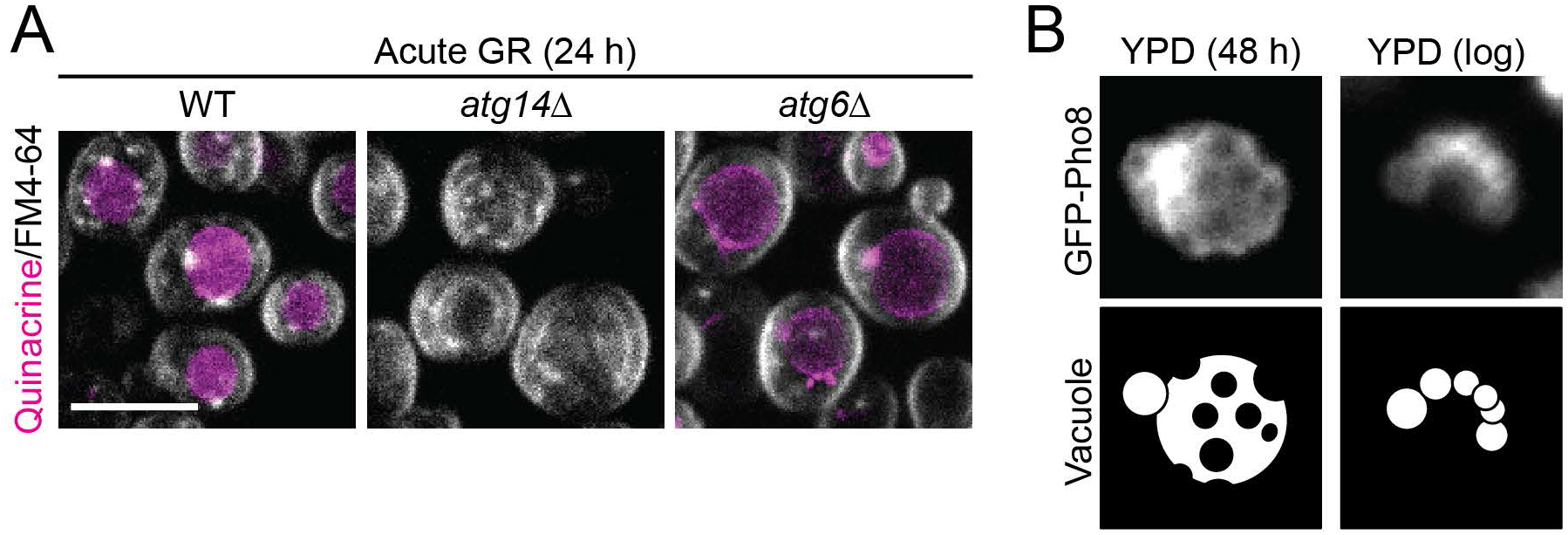
Vacuole phase partitioning by glucose restriction is associated with vacuole acidification, related to Figure 6. (A) Confocal images of cells pulsed by quinacrine and FM4-64 from indicated deletion strains. The image for WT (*left*) was duplicated from Figure 6D. Size bars, 10μm. (B) Epifluorescence images of cells expressing GFP-Pho8 after growing under rich media (YPD) either for 48 h or to the log phase without glucose starvation. Different vacuolar GFP-Pho8 localization patterns and shapes are illustrated in the bottom row.

**Movie S1. Different membrane characteristics resulted from defective ER-vacuole ergosterol exchange under glucose-restricted yeast vacuoles, related to Figure 2**

Time-lapse movies of *lam6Δ* cells expressing GFP-Pho8 under 40 h of glucose starvation. Maximum intensity projections of the vacuole labeled by GFP-Pho8 are shown in the top row. A single plane fluorescence image was generated along the yellow lines (x-z slice, *bottom*). L_o_, liquid-ordered membrane; L_d_, liquid-disordered membrane. Size bars, 1μm.

**Movie S2. Enhanced autophagosome production in cells under acute GR, related to Figure 3** Time-lapse movies of the control BY4742 strain expressing GFP-Atg8 after undergoing untreated (Fed) and 1 h of acute GR, gradual GR, or 250 nM rapamycin treatments. Sum intensity projections of each condition are presented. Size bars, 1μm.

**Movie S3. Selective L_o_ membrane internalization from the vacuole surface in glucose-restricted cells, related to Figure 5**

Time-lapse movies of the control W303 strain expressing GFP-Pho8 (cyan) and Ivy1-mCherry (magenta) after undergoing 24 h of acute GR. The equatorial focal plane from two different cells are presented. Size bars, 1μm.

